# A Dataset of Human Sperm Microscopy Videos for Detection, Tracking, and Motility Parameter Analysis

**DOI:** 10.64898/2025.12.15.694470

**Authors:** Atilla Sivri, JiWon Choi, Justin Bopp, Albert Anouna, Matthew VerMilyea, Gustave Alkhoury, Omer Onder Hocaoglu, Moshe Kam, Ludvik Alkhoury

## Abstract

We present HSMV, a dataset of Human Sperm Microscopy Videos acquired using two imaging systems. These were (i) a commercial Computer-Assisted Semen Analysis (CASA) system, and (ii) an optical microscope. The dataset contains samples from 51 healthy male participants and includes 3,161 video clips (approximately 26 hours of imagery). A subset of the clips from both systems was manually annotated with bounding boxes around each visible sperm head, linked across frames into trajectories, thereby establishing ground truth for detection, tracking, and motility analysis. Specifically, 14 CASA clips and 20 optical-microscope clips (each consisting of 900 frames) were manually annotated, yielding 29,900 frames and roughly 1.49 million sperm annotations. The dataset also includes thousands of unannotated clips captured under varying preparation conditions (washed vs. unwashed), dilution levels, magnifications, and regions of interest. When available, motility parameter reports are provided from either clinical technicians or CASA outputs. We demonstrate the use of this dataset in two example applications: (i) automated sperm detection and tracking using YOLOv5 and SORT, and (ii) visualizations of motility parameters. HSMV offers a comprehensive benchmark for developing and evaluating algorithms for sperm detection, tracking, classification, and motility assessment.

## Background and Summary

Semen analysis is a cornerstone of male fertility evaluation. It is often performed manually, by trained technicians who assess sperm concentration, motility, and morphology using an optical microscope [9]. In spite of following the guidance of the World Health Organization (WHO)[23], manual assessment suffers often from subjectivity and variability. These are the results of differences in operator skills and inconsistencies across laboratories[8, 23]. While Computer-Assisted Semen Analysis (CASA) systems have emerged to reduce subjectivity and improve reliability of fertility analysis, recent reviews call for additional efforts on validation and standardization [10, 3]. These would improve the accuracy and reproducibility of CASA results.

CASA systems leverage image processing algorithms and machine learning techniques to automatically detect, track, and analyze spermatozoa in video microscopy [25]. To advance such systems, there is a growing need for high quality, diverse, and richly annotated datasets that reflect the complexity of real-world laboratory conditions. Existing open-source datasets such as SVIA [5], SesViD [24], Hi-LabSpermTracking [1], VISEM [12], and VISEM-Tracking [20], contribute to this goal. Although very useful, each of these datasets comes with some limitations. VISEM includes no annotated frames, and consists of samples collected only from individuals with obesity. VISEM-Tracking expanded on VISEM by contributing approximately 650,000 bounding box annotations and object IDs to enable multi-object tracking, but again only from participants with obesity. SVIA includes approximately 66,000 bounding box annotations across 127 video clips, but doesn’t provide information about sample preparation conditions (e.g., chemical washing vs. raw unwashed, dilution, different zoom levels), and it used data collected only from CASA imaging systems. SeSViD provides 12 annotated videos for sperm detection, but provides short videos with relatively low frame rate. Finally, the Hi-LabSpermTracking dataset improves on prior work by supplying uninterrupted, long-duration videos with both bounding boxes and object IDs, but it also has low frame rate.

In this study, we build on these prior efforts by presenting a new dataset to complement and extend existing databases. It incorporates recordings from both an optical microscope and a CASA system, and includes manual annotations of sperm cells observed in the recordings. The dataset includes both manually annotated and unannotated video clips across the two imaging modalities, and several different sample preparation conditions (listed in the first paragraph of the Data Collection subsection of the Methods section). These additional annotations may enable a more representative benchmark for CASA systems while also supporting the development of new analytical methods that require rich datasets for training and testing.

Table 1 presents an overview of publicly available sperm video datasets, including their imaging systems, annotation volumes, and video counts. It situates our dataset within this landscape of available datasets for direct comparison.

**Table 1:** Comparison of public sperm video datasets, including microscopy systems, preparation conditions, annotation volume, and video availability.

| Dataset | Microscopy Systems | Preparation Conditions | # of Annotations | # of Videos |
| --- | --- | --- | --- | --- |
| SVIA | CASA | Unwashed, 640x480, 30 fps | 125,000 | 101 annotated (1-2 seconds) |
| SeSViD | Optical Microscope | Unwashed, 640x640, 5 fps | 92,329 | 12 annotated (2-30 seconds) |
| Hi-LabSperm Tracking | Smartphone and Optical Microscope | Unwashed, 1920x1080, 10 fps | 727,293 | 66 annotated (60 seconds 10 fps) |
| VISEM | Optical Microscope | Unwashed, 640x480, 30 fps | 0 | 336 unannotated (30 seconds) |
| VISEM Tracking | Optical Microscope | Unwashed, 640x480, 30 fps | 600,000 | 20 annotated (30 seconds) |
| HSMV (this work) | CASA and Optical Microscope | Washed, unwashed, 100-400x magnification levels, different dilution levels 1280x1024-1320x1080 30-59.94 fps | 1,490,875 | 34 annotated (14 CASA and 20 optical microscope) and 3,127 unannotated (30 seconds) |

To our knowledge, HSMV is the first publicly available dataset to provide manually annotated sperm recordings from both CASA and optical microscope modalities, which are also paired with motility parameters metadata. Our dataset may thus be useful in the development and evaluation of next-generation CASA systems. It includes data collected from the following: (1) participants drawn from the general population, rather than from a specific clinical population; (2) multiple imaging modalities, specifically, images captured using both an optical microscope and a CASA system; (3) a range of sample preparation methods with robust manual annotations; and (4) comprehensive motility information. This dataset thus provides a foundation for research in object detection, tracking, and automatic motility parameter calculation and analysis, and may contribute toward more accurate, reproducible, and generalizable automated semen analysis workflows.

All codes used in this paper are available at https://github.com/DFL-KamLab/HSMV_A-Dataset-of-Human-Sperm-Microscopy-Videos and all videos, bounding box annotations, and motility data are found at: https://huggingface.co/datasets/DFL-KamLab/HSMV_A-Dataset-of-Human-Sperm-Microscopy-Videos

## Methods

### Data Collection

The recordings in our dataset were collected at two different sites: the Sperm and Embryo Bank of New Jersey in Mountainside, New Jersey, using an optical microscope (see the Optical Microscope Recordings subsection), and the Ovation-Fertility Laboratory in Brentwood, Tennessee, using a CASA system (see the CASA Recordings subsection). Semen samples were captured under two magnification levels (100x and 200x for the optical microscope, 200x and 400x for the CASA system). Samples were prepared under the following conditions: (a) raw, unwashed samples, that is, freshly collected semen with no further treatment or processing; (b) chemically washed samples, prepared by removing the seminal plasma and replacing it with a buffer medium, a standard laboratory step to isolate motile spermatozoa; and (c) diluted samples, prepared for a small number of participants by adding a buffer where the concentration was too high to be imaged reliably. The dilution process happened at a ratio of 1-to-5 for the optical microscope recordings, and at 1-to-5 and 1-to-10 for the CASA recordings. Samples were allowed to liquefy for 10 to 20 minutes at 37^◦^C before preparation. Washed and unwashed preparations were produced by each clinic as part of its routine procedure, following the sperm preparation methods described in the sixth edition of the WHO laboratory manual for the examination and processing of human semen [23]. The procedures used at each site are given below, in the Optical Microscope Recordings and CASA Recordings subsections.

### Optical Microscope Recordings (Swift M10DB-MP Series)

The optical microscope recordings were collected at the Sperm and Embryo Bank of New Jersey using a Swift M10DB-MP Series Advanced Microscope [13] equipped with a Fujifilm X-T10 camera [7]. Video clips were acquired at a resolution of 1920x1080 pixels and a frame rate of 59.94 fps. These recordings exhibit variability in magnification levels (100x and 200x), with approximately equal representation of each level, as well as balanced distribution of sample preparation conditions (chemically washed, or unwashed raw). Washed samples were prepared by density gradient centrifugation, beginning 30 to 60 minutes after collection. A gradient was formed from 2.0 mL of the 80% PureCeption solution (Sage) with the 40% solution layered on top, and up to 1.5 mL of semen was layered above it. The tube was centrifuged at 425 g for 20 minutes, the pellet was transferred to a fresh tube, and 4.0 mL of Modified Sperm Washing Medium (modified HTF with 5.0 mg/mL bovine albumin, Irvine Scientific) was added. After rotating for 5 minutes and a second centrifugation at 425 g for 8 minutes, the supernatant was removed to the 0.6 mL mark and the pellet was resuspended. For the optical microscope recordings, we extracted calibration factors for both magnification settings (100x and 200x) to convert the pixel-based measurements into microns (*µ*m). The calibration factors were obtained by measuring a prefixed physical dimension on a calibration slide and mapping the corresponding number of pixels to their equivalent length in *µ*m. The calibration slide, the measurements, and the resulting images for this system are given respectively in the Calibration appendix, in Figure A1, Table A1, and Figure A2. Throughout this paper, we use calibration factors of 0.419 *µ*m/px and 0.202 *µ*m/px for the 100× and 200× magnification settings, respectively. In total, recordings of semen samples were collected from 24 healthy participants. We divided the original recordings into 30-second clips, resulting in a total of 3,021 clips, and cropped into a resolution of 1320x1080 pixels. Each clip typically captures a specific region within the microscope’s field of view, which we refer to as a *scene*. We show in Figure 1 a schematic of the extraction of clips from a recorded scene, magnification differences, unwashed and washed samples, and dilution differences.

**Figure 1:**
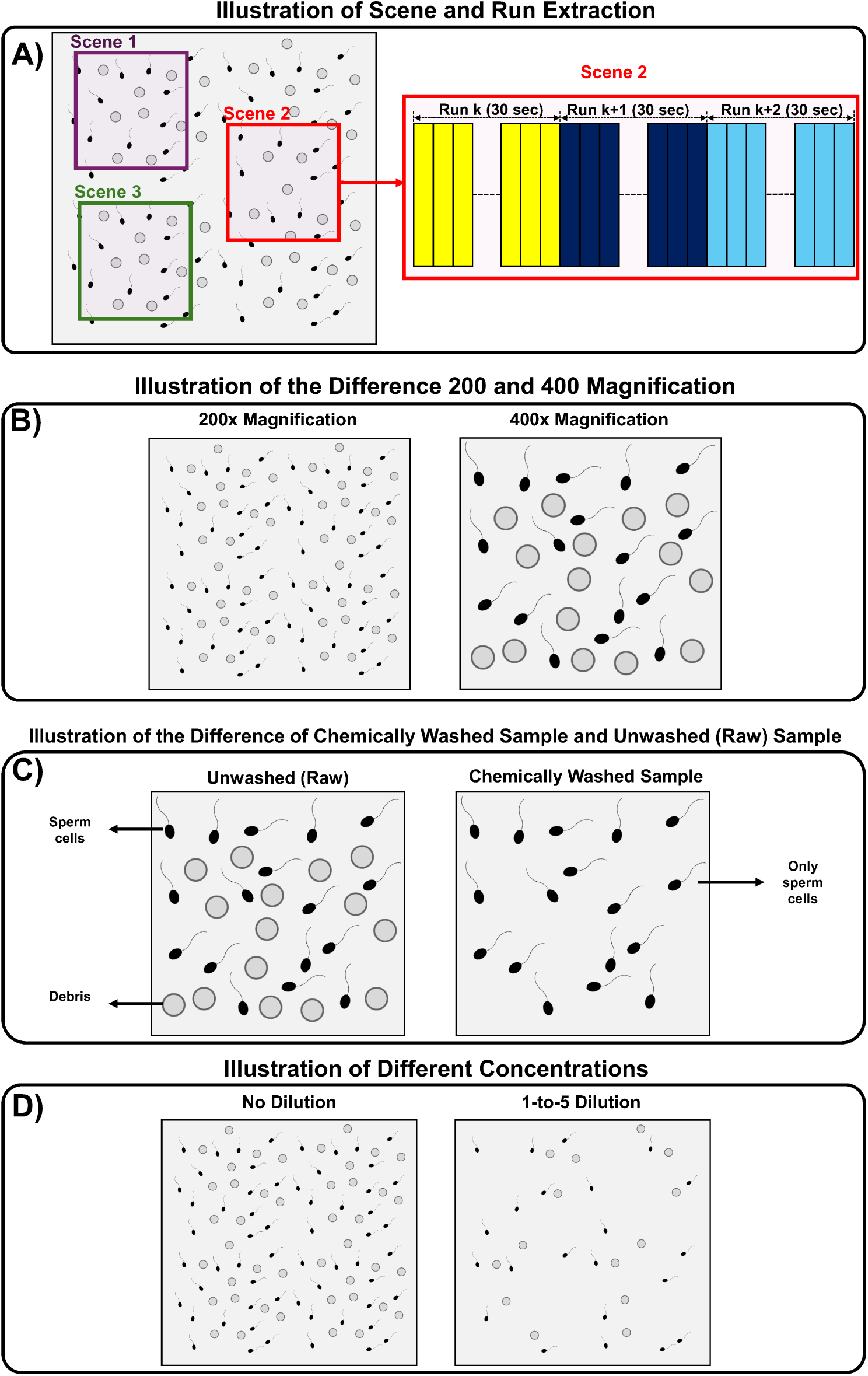
A) Illustration of the scene and run extraction process. Representative regions of the full field of view are selected, each of which is a scene, and each scene is recorded and divided into multiple 30-second runs, also referred to as clips, to create consistent video samples for annotation and analysis. B) Illustration showing the difference between two magnification levels. C) Illustration showing the difference between unwashed (raw) and chemically washed samples. D) Illustration showing the difference between a sample with no dilution and with 1-to-5 dilution.

Out of the samples of 24 participants, 20 samples were chosen to be manually annotated, with one clip per participant partially manually annotated (900 frames each). In addition to the video data, laboratory technicians provided a sperm count and concentration report for both chemically washed and raw unwashed samples. These reports include measures such as sample volume (mL), motile and non-motile counts (in millions), concentration (million/mL), total cells (million/mL), percentage motility, and forward motility index. The full reports are given in Tables A10 and A11 of the Dataset Property Tables appendix.

### CASA Recordings from Sperm Class Analyzer, SCA

The CASA recordings were collected at the Ovation-Fertility Laboratory (Brentwood, TN, USA) using a Sperm Class Analyzer (SCA) system[19] manufactured by MICROP-TIC S.L. Video clips were acquired at a resolution of 1280x1024 pixels and a frame rate of 30 fps. The video clips were originally recorded as 30-second clips, so no further partitioning was necessary.Washed samples were prepared by density gradient centrifugation. A gradient was formed from 1.5 mL of the lower layer and 1.5 mL of the upper layer of ISolate (Irvine Scientific, 99264), and up to 4 mL of semen was layered on top. The column was centrifuged at 1,500 rpm for 15 minutes, the medium above the pellet was removed, and the pellet was resuspended in 4 to 6 mL of Multipurpose Handling Medium Complete (MHM-C, Irvine Scientific, 90166). After a second centrifugation at 1,500 rpm for 5 minutes, the supernatant was discarded and the pellet was resuspended in 0.5 mL of MHM-C. These recordings were acquired at two magnification levels (200x and 400x), with approximately equal representation of each, as well as balanced distribution of sample preparation conditions (chemically washed, or unwashed raw), and dilution levels (no dilution, 1 to 5, 1 to 10).

Similar to the optical microscope, we extracted calibration factors for both magnification settings (200x and 400x) for the CASA recordings. The stage micrometer, the measurements, and the resulting images for this system are given in the Calibration appendix, in Figure A3, Table A2, and Figure A4 respectively. Throughout this paper, we use calibration factors of 0.397 *µ*m/px and 0.198 *µ*m/px for the 200× and 400× magnification settings, respectively.

In total, semen samples were collected from 27 participants, yielding a total of 140 30-second-long clips.

Out of the samples of the 27 participants, samples of 14 of them were chosen at random to be manually annotated, with one clip per participant partially manually annotated (900 frames each). Alongside the video clips, the CASA system automatically generated motility parameter reports for each participant, including sample volume (mL), percentage of rapid, slow, non-progressive, and immotile sperm, and sperm concentration (million/mL). The full reports are given in Tables A8 and A9 of the Dataset Property Tables appendix.

### Ethics, Recruitment, and Informed Consent

Optical microscope images (SEBNJ). Participants were recruited through an advertisement on the sperm bank’s website and by informing individuals who contacted the bank regarding donation. A short telephone screening established eligibility: male, aged 18 to 35, and no participation in the campaign during the preceding 12 months. Beyond age, sex, and the date of any previous donation, no personal or clinical information was recorded. Participants signed informed consent upon arrival and before any sample was provided, covering both participation and the public release of the recordings, and received monetary compensation.

To protect participant information, each participant was given a tag bearing a randomly generated identifier, used only to label the specimen and surrendered before departure; the identifier was never recorded alongside the participant’s name. The donor list was held as a single secured copy at the sperm bank, was not shared with the research team, and names were deleted within 12 months of collection, or immediately upon preparation of any document intended for external sharing. No physical samples were transferred, only the recordings and the accompanying laboratory measurements. All procedures were approved by the Institutional Review Board of the New Jersey Institute of Technology under protocol 2106008579.

CASA images (Ovation Fertility). Recordings were obtained from patients of the Ovation Fertility Laboratory under the clinic’s standard procedures, which include written consent permitting de-identified data to be used for research and publication. Recruitment, sample handling, recording, and de-identification were carried out by the clinic; the research team received no physical samples, only the de-identified recordings and the motility reports generated by the clinic’s CASA system. All procedures for this data transfer were approved by the Institutional Review Board of the New Jersey Institute of Technology under protocol 2110013259.

### Data Processing

For consistency, all video clips were standardized into 30-second clips with a unified naming convention, described in the Data Records section. Since the outer edges of the images were blurred, each optical microscope recording was cropped to 1320x1080 pixels and segmented into consecutive 30-second clips using MATLAB. The CASA video clips were already supplied in the desired 30-second format.

Both recordings consist primarily of unannotated clips. The manually annotated portion is smaller but carefully reviewed, yielding high-quality ground truth. In the optical microscope recordings, 20 clips (1 clip per subject) of 900 frames each (except one video clip, annotated up to 200 frames) were manually annotated, while in the CASA recordings, 14 clips (1 clip per subject) of 900 frames each were manually annotated.

Annotation of sperm heads was performed by the research team using Labelbox (https://labelbox.com/) for the optical microscope recordings and using a MATLAB-based annotation script for the CASA recordings. For non-motile sperm cells of the optical microscope recording, we used Labelbox’s auto-propagation tool, which automatically replicates bounding boxes across frames. This feature prevents the need to manually annotate cells that maintain a constant position throughout the recording. Annotations were cross-verified by multiple team members. Ground truth annotations for each sperm cell are provided in the following format: i cx cy w h track id. A detailed description of each one of the annotation parameters is presented in Table 2.

**Table 2:** Bounding-box and trajectory annotation format and description for sperm cell annotations.

| Parameter | Description | Range | Notes |
| --- | --- | --- | --- |
| *i* | Class index | 0 | Only one class (sperm cell) |
| *cx* | Normalized x-coordinate of the bounding box center | 0-1 | Relative to frame width |
| *cy* | Normalized y-coordinate of the bounding box center | 0-1 | Relative to frame height |
| *w* | Normalized bounding-box width | 0-1 | Fraction of frame width |
| *h* | Normalized bounding-box height | 0-1 | Fraction of frame height |
| *track_id* | Trajectory identity | Integer | Same value across frames marks the same sperm cell; unique within a frame |

For example, the annotation 0 0.45 0.62 0.08 0.12 7 denotes a sperm head centered at 45% of the frame width and 62% of the frame height, with a bounding box spanning 8% of the width and 12% of the height, belonging to trajectory 7.

In addition to pre-processing and annotation, we also applied automated detection and tracking methods to illustrate potential use-cases of the dataset. Specifically, YOLOv5[14] was employed for object detection, and SORT[4] was used to generate trajectories from both ground truth and YOLOv5 detections. From the tracks generated by SORT, motility parameter calculations were performed. These methods, along with their results, are described in detail in the *Detection and Tracking Experiments* appendix, in the subsections *Object Detection using YOLOv5*, *Tracking using Simple Online and Realtime Tracking (SORT)*, and *Motility Parameters*.

In addition to the manually annotated images, the dataset repository contains Python scripts used to run YOLOv5 and SORT, and to compute motility parameters. These are provided to facilitate easy reproducibility.

### Data Records

The complete dataset is deposited on the Hugging Face Hub [18] and is distributed under a Creative Commons Attribution 4.0 International license (CC BY 4.0). It comprises the video recordings, a metadata file for each recording, manually annotated sperm cell bounding boxes for a subset of the recordings, and the motility reports produced by the laboratory technicians or by the CASA system.

The dataset is organized following a structure inspired by the Brain Imaging Data Structure (BIDS) standard [11]. BIDS is a community convention for naming and arranging neuroimaging data that provides systematic organization and enhanced reproducibility. As in BIDS, the directory hierarchy of our system is mirrored in the name of each file. The top level is the acquisition system, ‘sys-opt’ for the optical microscope and ‘sys-casa’ for the CASA system. The next level is the type of data stored; unprocessed recordings are held under ‘rawdata’. Below that come the participant identifier and then the session number, where a session denotes one visit for data collection, so that a participant returning on a later date is recorded as a second session. Within a session, a *scene* is a region of the microscope’s field of view that was recorded once, and a *run* is a single 30-second video taken from that scene, as illustrated in Figure 1. Runs are numbered sequentially across the whole session, and the metadata of each run records the scene it came from. The terms *run* and *clip* are used interchangeably throughout this paper. Because the two systems differ in frame rate, a clip holds 900 frames on the CASA system and 1,800 frames on the optical microscope. Ground truth annotations are stored alongside the run they belong to.

Tables 3 and 4 give an overview of the dataset for both acquisition systems. Table 3 covers the dataset as a whole, and Table 4 the manually annotated subset, reporting for each system the number of participants, the number of clips, and the total recording time.

**Table 3:** Summary of the complete dataset, manually annotated and unannotated clips combined. For each recording system, we report the number of participants, number of clips, and total video length in seconds.

| **Recording System** | **Participant count** | **Clip count** | **Total Length (sec)** |
| --- | --- | --- | --- |
| Optical Microscope | 24 | 3,021 | 90,630 |
| CASA | 27 | 140 | 4,200 |

**Table 4:** Manually annotated subset summary. For each recording system, we report the number of participants, number of clips, total video length in seconds, and overall number of annotated sperm cells.

| Recording System | Participant count | Clip count | Total Length (sec) | # of annotations |
| --- | --- | --- | --- | --- |
| Optical Microscope | 20 | 20 | 300 | 802,677 |
| CASA | 14 | 14 | 420 | 688,198 |

The CASA recordings came from 27 participants, with manually annotated bounding boxes available for 14 of them. For each of these 14 participants we annotated one complete clip, that is, all 900 frames of a 30-second recording acquired at 30 fps.

The optical microscope recordings came from 24 participants, with manually annotated bounding boxes available for 20 of them. For each of these 20 participants we annotated one clip in part rather than in full: the first 900 frames, which at 59.94 fps corresponds to the first 15 seconds of the recording. One clip was annotated to 200 frames only.

This structure provides flexibility for multiple research directions; the unannotated videos can support unsupervised and semi-supervised learning approaches, while the manually annotated subset offers verified ground-truth data suitable for evaluating supervised methods for sperm detection and tracking.

Each acquisition system (optical microscope and CASA systems) occupies a top-level directory, sys-casa or sys-opt. Within each, unprocessed raw recordings are stored under rawdata by participant and then by session. A session directory holds one *.avi* recording and one *.json* metadata file per run, sharing a common base name. Where a run belongs to the manually annotated subset, a sibling directory with the suffix gt holds its annotations, one plain-text file per frame, indexed from zero. Motility reports are held at the top level of each system directory.

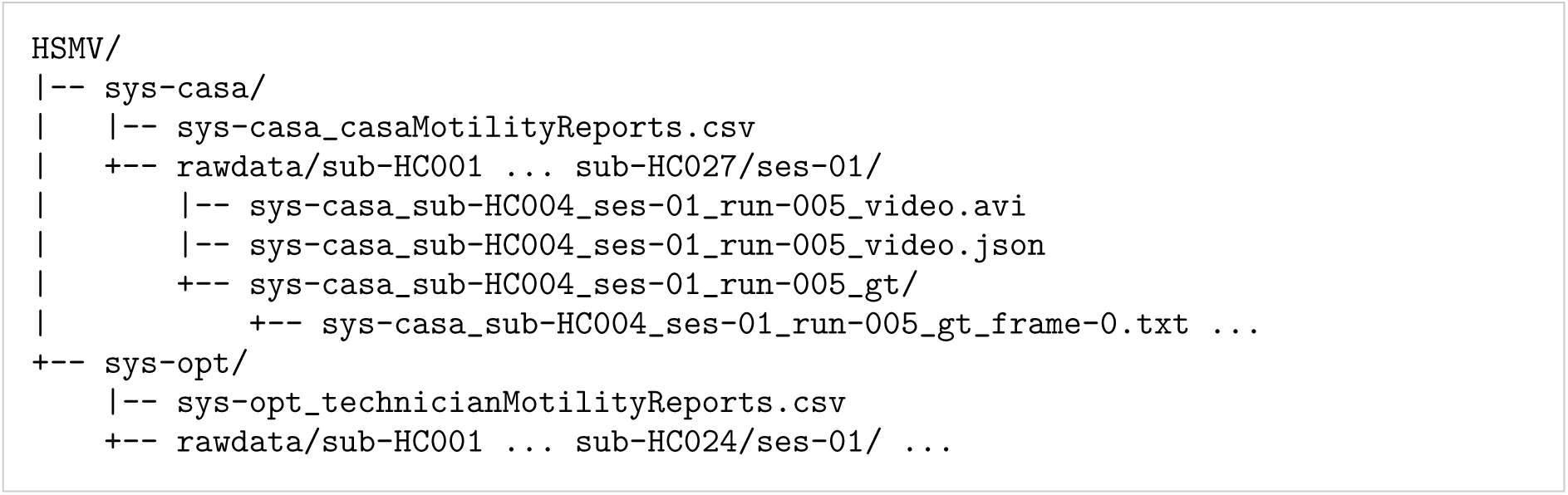

Each file name encodes critical information about the system used for collection (CASA or optical microscope), subject ID, session number, and the run number. Sessions represent data collections that happen on a given day. Runs are numbered sequentially within a session and span all of its scenes, so two runs from the same session may differ in scene, preparation, magnification, or dilution.

For example, the file name:

sys-casa sub-HC004 ses-01 run-005 video.avi indicates that the AVI video clip originates from the CASA dataset, came from subject HC004, was captured during session 01, and represents the fifth run.

In addition to the *.avi* recording named above, each recording is paired with a *.json* file (for this example, the file name is: sys-casa sub-HC004 ses-01 run-005 video.json) that captures all essential metadata. The file includes experiment-level details such as the experimenter, institution, department, device manufacturer and model, software versions, and a brief description of the procedure. It also records the technical and recording-specific parameters, such as sampling frequency, magnification, dilution ratio, sample type (washed or unwashed), system type (e.g., “casa”), subject ID, session number, run number, scene number, and the date of acquisition. The complete *.json* file for this recording is given in Listing 1.

For each video frame, we generate one text file holding the ground truth annotations of that frame. The file name repeats the name of the video and appends the frame number, indexed from zero, so that the first frame of sys-casa sub-HC004 ses-01 run-005 video.avi is annotated in sys-casa sub-HC004 ses-01 run-005 gt frame-0.txt. Each line of the file describes one sperm cell as a bounding box in the normalized YOLO format i cx cy w h, followed by its trajectory identity track id, as set out in Table 2. Figure 2 shows a video frame containing manually annotated sperm cells, recorded with the CASA system in panel (a) and with the optical microscope in panel (b), with the green boxes marking the annotated sperm heads.

**Figure 2:**
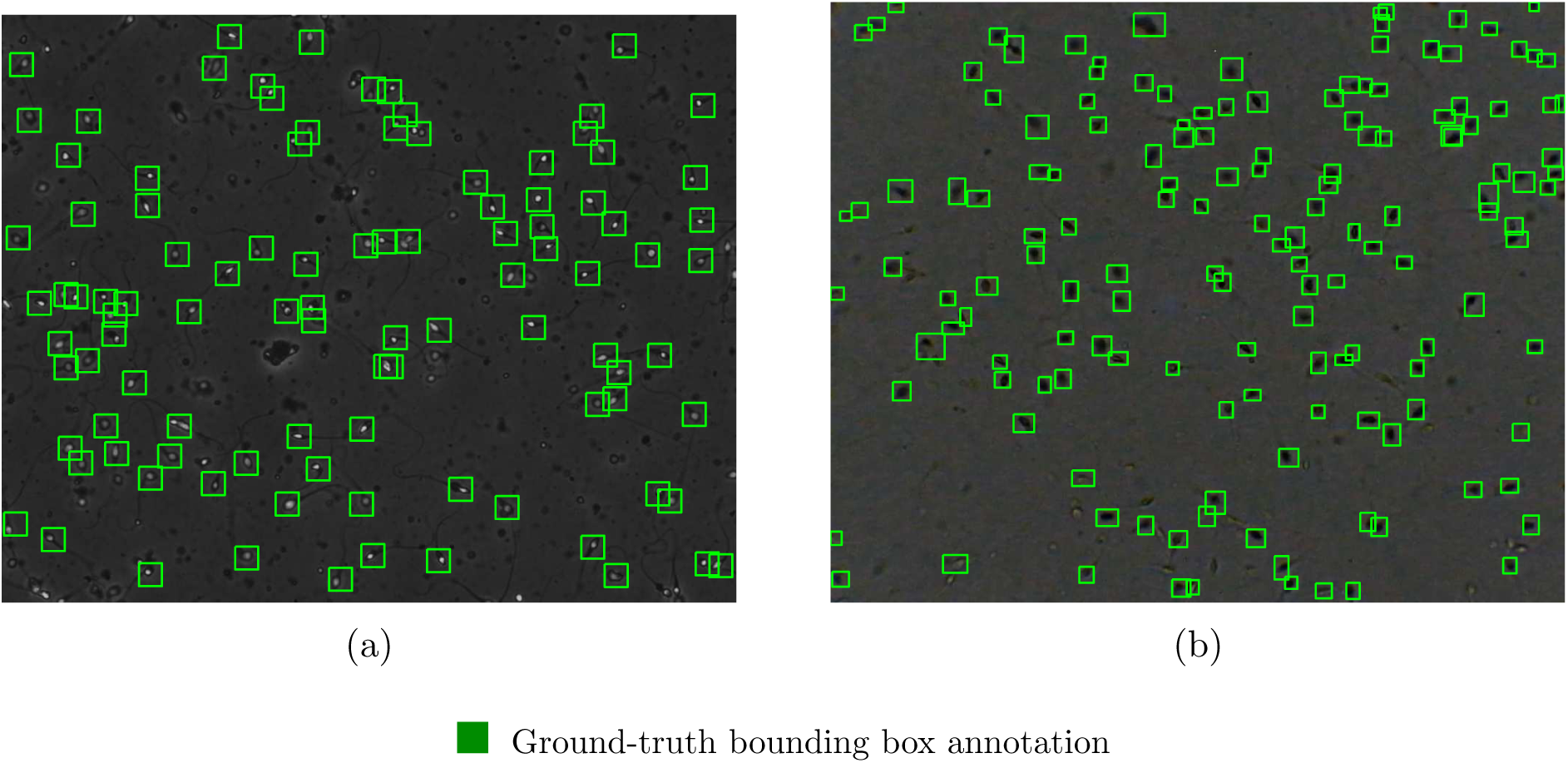
Ground-truth annotations provided with the dataset, shown for one frame from each acquisition system. (a) Frame 50 of sys-casa sub-HC004 ses-01 run-005, recorded with the CASA system. (b) Frame 50 of sys-opt sub-HC010 ses-01 run-049, recorded with the optical microscope. Green boxes mark the manually annotated sperm heads that constitute the ground truth. Each box corresponds to one line of the associated per-frame annotation file, given in normalized YOLO coordinates as set out in Table 2.

In addition to the recorded videos, we provide two motility reports per session, one for the washed sample and one for the unwashed sample. For the CASA recordings, the reports were generated by the Sperm Class Analyzer and provide the sample volume (mL), the percentages of rapid, slow, non-progressive, and immotile spermatozoa (%), the sperm concentration (10^6^/mL), and the total count (10^6^). For the optical microscope recordings, the reports were produced by a laboratory technician, separately for the unwashed and washed samples, and provide the sample volume (mL), the motile and non-motile counts (10^6^/mL), the concentration (10^6^/mL), the total cells (10^6^), the percentage motility (%), and a forward motility index graded from lowest score of 0 to highest score of 3.

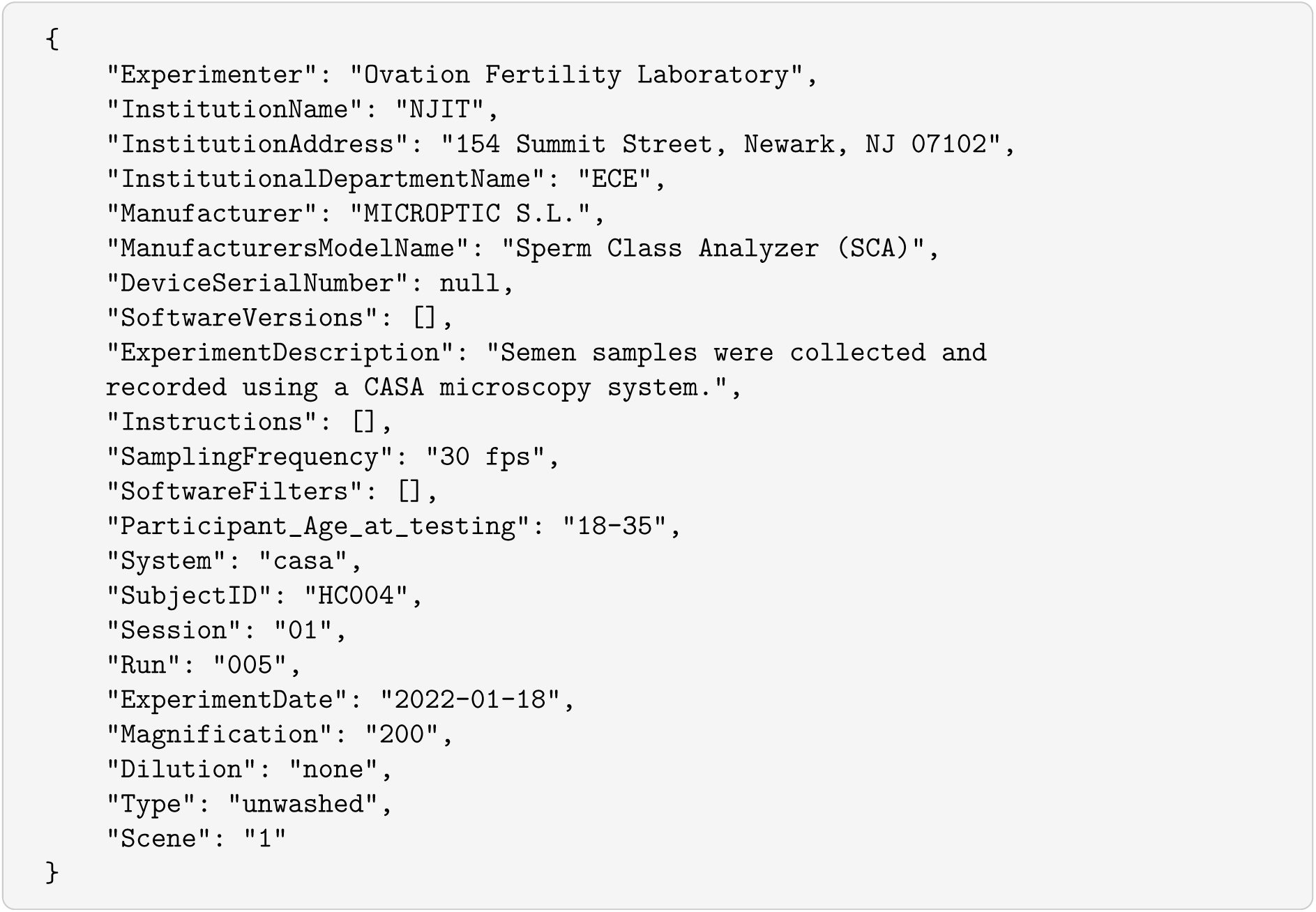

Listing 1: Metadata file accompanying a single recording, sys-casa sub-HC004 ses-01 run-005 video.json. Every property that distinguishes one recording from another is recorded here.

The BIDS-like structure of the dataset, together with the metadata file that accompanies every recording, allows a user to seamlessly gather recordings of a given specification without opening the videos themselves. Every property that distinguishes one recording from another, such as the acquisition system, the preparation type, the magnification, and the dilution, is recorded in the metadata; a representative example of the full set of entries is given in Listing 1. For example, a user interested in the optical microscope recordings that are washed and acquired at 200× can run a short script that reads the metadata files, selects those whose fields match, and returns the corresponding videos and annotations.

The dataset is maintained as a versioned resource; the version described in this paper is v1.1. We intend to continue extending the resource, and subsequent releases may add further recordings and further annotations. Each release is issued as a new version with its own identifier, and readers are encouraged to consult the repository, which always lists the most recent version and the changes it contains.

By embedding essential information directly within the file names, providing metadata, and maintaining a hierarchical directory structure, the system supports easy implementation and streamlines the integration of new data.

The dataset properties are given in full in the Dataset Property Tables appendix: the clip counts per participant in Tables A4 and A5, the annotated sperm cells and annotated frames per participant in Tables A6 and A7, and the motility parameters in Tables A8 to A11. The tables also list the motility parameters calculated by the CASA system, including volume, rapid %, slow %, non-progressive %, immotile count, and sperm concentration. They further list the parameters obtained by laboratory technicians using a microscope, including volume, motile count, non-motile count, concentration, total cells, motility %, and forward motility index.

### Technical Validation

This section describes the quality control applied to the dataset, and then demonstrates three uses the dataset supports, namely: sperm detection, multi-object tracking, and the computation of kinematic parameters. The models used and the settings under which they were run are given in the Detection and Tracking Experiments appendix, together with the detection results across five YOLOv5 variants in Table A3, the detections and tracks for both systems in Figures A9 and A10, and the kinematic parameters computed from ground truth and from YOLOv5 detections in Figures A11 to A14.

#### Quality control

Every recording and every annotation in the dataset passed through the following procedure:

1. Recordings were acquired and de-identified at the collecting laboratory.
2. Each recording was divided into 30-second clips, and its file name, contents, and metadata were checked against the conditions under which it was acquired.
3. Each manually annotated clip, including the trajectory identities, was annotated by one annotator and reviewed frame by frame by a second, who corrected any errors. Where the two disagreed, they discussed the case and agreed how to proceed. The reconciled annotations are the ground truth released with the dataset.
4. The complete dataset was verified locally, then uploaded to the repository and verified again there.

#### Detection

We trained YOLOv5 [14] on the manually annotated optical microscope subset, splitting the data 80/20 by video so that no recording contributes to both the training and the testing set. On the held-out testing set the model reached a precision of 0.66 and a recall of 0.76, giving an F1-score of 0.71. For the CASA recordings we did not train a model of our own; we applied the weights released with the VISEM-Tracking dataset [20], which were trained on optical microscope recordings, to all 14 manually annotated CASA subjects. This gave a precision of 0.82 and a recall of 0.43, and an F1-score of 0.56.

Tracking and kinematic parameters. We used the SORT algorithm [4] to generate sperm trajectories, (a) from the ground truth annotations and (b) from the YOLOv5 detections. From each set of trajectories we computed eight kinematic parameters in common clinical use, comprising three velocities (VCL, VSL, VAP), three dimensionless ratios describing the shape of the path (LIN, WOB, STR), and two measures of lateral and angular deviation (ALH, MAD). Velocities and displacements are expressed in physical units using the micron-per-pixel ratios reported in Table 5. Figure 3 summarizes these parameters for representative participants from each system, computed from ground truth trajectories and from YOLOv5 detections.

**Figure 3:**
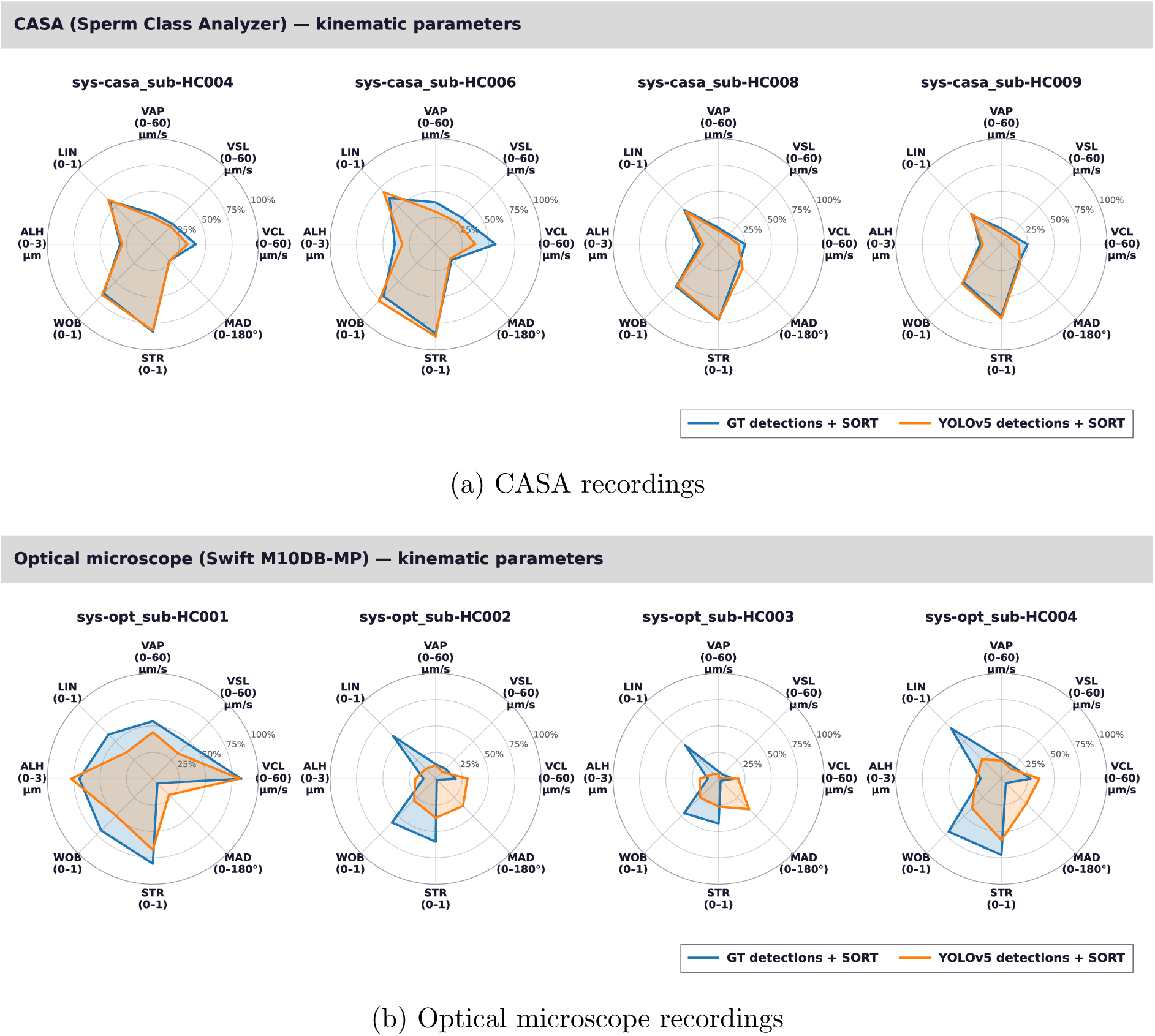
Kinematic parameters for four representative participants from each acquisition system, shown as spider plots. Within each panel, the blue trace gives the values obtained from the manually annotated ground truth trajectories and the orange trace those obtained from the YOLOv5 detections, across the eight kinematic parameters. (a) CASA recordings, acquired at 30 fps with a micron-per-pixel ratio of 0.397 or 0.198 depending on magnification. (b) Optical microscope recordings, acquired at 59.94 fps with a ratio of 0.419 or 0.202. The two systems therefore differ in temporal and spatial resolution, and the panels are not intended for direct numerical comparison between systems; each shows the agreement between annotation-derived and detection-derived parameters within its own system.

**Table 5:**
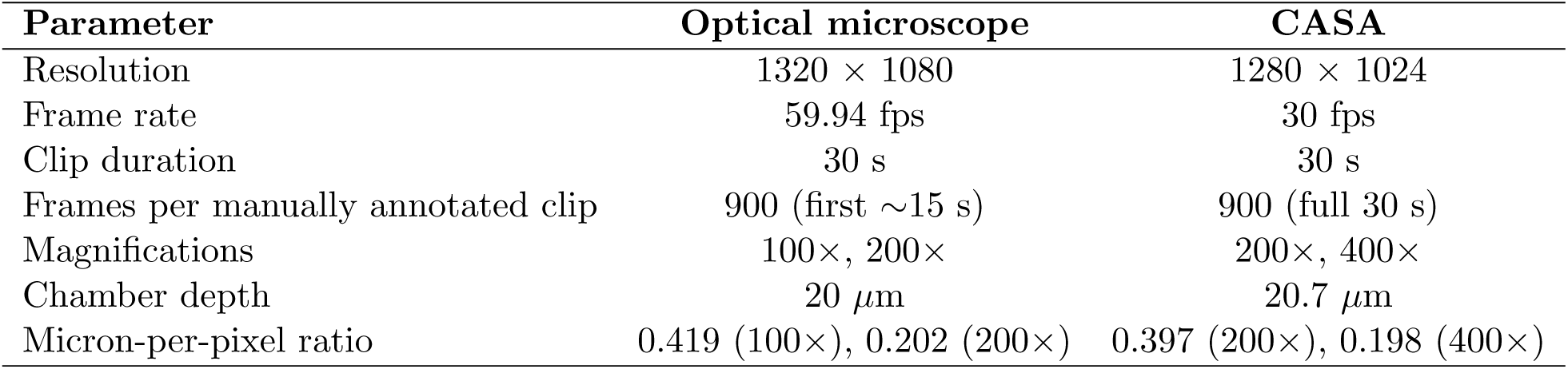
Acquisition parameters for the two systems. The magnification each micronper-pixel ratio corresponds to is given in parentheses.

| Parameter | Optical microscope | CASA |
| --- | --- | --- |
| Resolution | 1320 × 1080 | 1280 × 1024 |
| Frame rate | 59.94 fps | 30 fps |
| Clip duration | 30 s | 30 s |
| Frames per manually annotated clip | 900 (first ~15 s) | 900 (full 30 s) |
| Magnifications | 100×, 200× | 200×, 400× |
| Chamber depth | 20 $\mu\text{m}$ | 20.7 $\mu\text{m}$ |
| Micron-per-pixel ratio | 0.419 (100×), 0.202 (200×) | 0.397 (200×), 0.198 (400×) |

### Usage Notes

All video clips in this dataset are provided in AVI format. The manually annotated subsets include bounding boxes for spermatozoa, enabling precise evaluation of detection and tracking algorithms. Together, the manually annotated and unannotated material offer a versatile resource for both developing new computer vision methods and supporting applications such as motility analysis and training of laboratory personnel.

The dataset supports both supervised and unsupervised learning workflows. For supervised training, the manually annotated subsets can be used directly to benchmark sperm detection algorithms. For semi-supervised or self-training approaches, models trained on the manually annotated data can generate pseudo-labels for the larger unannotated video clips, which can then be refined by domain experts to accelerate and scale annotation.

Beyond detection, the manually annotated tracks enable training and validation of sperm tracking methods, which in turn support motility analysis. Motility parameters such as velocity, linearity, and curvilinear distance can be computed either from ground truth annotations or from automated detections (see the Motility Parameters section of the Detection and Tracking Experiments appendix). These parameters can be summarized in clear, interpretable reports, allowing non-technical users to extract clinically relevant insights and compare them with technician assessments.

In addition, the dataset can serve as a valuable training resource for technicians. By comparing their own motility measurements with values generated by the CASA system and those obtained through manual evaluation, technicians can calibrate their assessments, improve accuracy, and enhance consistency in clinical practice.

This dataset is our initiative to build a large, richly annotated resource that spans multiple imaging systems and pairs the recordings with motility reports. It remains a work in progress: further recordings, annotations, and analysis representations may be added in future releases, and we encourage users to consult the dataset repository [18] for updates.

The manual annotations released with HSMV establish ground truth by human judgment, and are therefore constrained by the expertise of the annotators who produced them. For work that requires ground truth known exactly rather than by agreement, the simulation tool of Choi et al. [6] can generate synthetic sperm image sequences from configurable parameters, including the number of cells, their concentration, and their trajectories, and can be used alongside these recordings.

We also provide pycasa [17], an open-source Python toolkit that takes a semen video as input and produces detections (YOLOv5 [14], digital washing [2], and the method of Urbano et al. [21]), tracks (SORT [4], DeepSORT [22], and JPDAF [16]), motility parameters (kinematic and those defined in the WHO laboratory manual [23]), and visualizations. Several methods are available at each step, so pipelines can be compared on the same recordings. It ships with a recording from this dataset, so it can be run without any local data collection, and is available under an Apache 2.0 license at https://github.com/DFL-KamLab/pycasa.

## Data Availability

The dataset described in this work is available on the Hugging Face Hub[18] at https://huggingface.co/datasets/DFL-KamLab/HSMV_A-Dataset-of-Human-Sperm-Microscopy-Videos

It is distributed under a Creative Commons Attribution 4.0 International licence (CC BY 4.0) and comprises the video recordings, their metadata, the manual annotations, and the motility reports.

## Code Availability

All of the codes used in this paper could be found on the following GitHub repository: https://github.com/DFL-KamLab/HSMV_A-Dataset-of-Human-Sperm-Microscopy-Videos.

## Acknowledgments

We thank the participants who consented to provide samples for this study. We are grateful to the clinical staff at the Sperm and Embryo Bank of New Jersey (SEBNJ) and Ovation-Fertility Laboratory in Brentwood, Tennessee, for their assistance in data collection and for providing technician-reported motility parameters. We also acknowledge the contributions of the research assistants Omer Hakan Yaran, Asmar Hasanova, Dorukhan Cakir, John Holck, and Hasan Dagdelen in Data Fusion laboratory at the New Jersey Institute of Technology for their support in annotation and preprocessing. Finally, we thank Muhammet Tayfur Ertas, Istanbul Medipol University, for a helpful exchange regarding the calibration of the two imaging systems. This work was conducted under IRB approvals 2106008579 and 2110013259.

## Author Contributions

A.S., J.C., M.K, and L.A. conceived and designed the study. A.S., J.C., J.B., A.A., M.V. and L.A. collected the data. A.S., J.C. and L.A. processed and standardized the data. A.S., J.C., G.A., O.O.H. and L.A. performed the annotations. A.S. and L.A. conducted the deep learning experiments and computed motility parameters. A.S. and L.A. validated and visualized the results. A.S., G.A., O.O.H. and L.A. wrote the manuscript. M.K. and L.A. supervised the project. All authors reviewed and approved the final manuscript.

## Funding

Not applicable.

## Competing Interests

L.A. holds equity in and serves as a founding engineer for Cognitive Signals Inc. These interests have been disclosed to Weill Cornell Medicine and are managed in accordance with its conflict-of-interest policies. The remaining authors declare no competing interests.

## APPENDIX

Supporting calibration procedures and dataset property tables

### Overview

This appendix accompanies the Human Sperm Microscopy Videos (HSMV) dataset and the analyses presented in the main text. It presents detailed spatial calibration procedures for both the optical microscope and CASA imaging systems, including theoretical and empirically measured micron-per-pixel ratios. In addition, it provides specimen-level tables summarizing video counts, annotation statistics, and motility data for the recordings included in the study.

The calibration section details both theoretical and empirically measured micron-per-pixel ratios for the optical microscope and CASA imaging systems. Calibration measurements are supported by image-based calibration targets and representative sperm head examples. In summary, the calibration factors we use in the main text are extracted using the empirical method. For completeness, we also report theoretically derived calibration values in the Theoretical calibration method section; however we recommend adopting the empirically measured micron-per-pixel ratios summarized below, and these ratios were used in the main text.

Summary of empirically measured micron-per-pixel ratios:

1. Optical microscope system (empirical ratios, see the Optical Microscope section below for more details):

- 100x magnification → micron-per-pixels ratio = 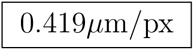
- 200x magnification → micron-per-pixels ratio = 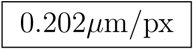
2. CASA system (empirical ratios, see the CASA system section below for more details):

- 200x magnification → micron-per-pixels ratio = 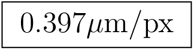
- 400x magnification → micron-per-pixels ratio = 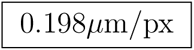

In addition, the tables presented in the Dataset Property Tables appendix provide detailed specimen-level information supporting the analyses described in the HSMV study. These tables include video clip counts; annotation counts; the total number of manually annotated and unannotated frames; and comprehensive motility parameters for both CASA and optical microscope recordings, along with sample metadata such as collection dates, specimen volumes, motility classifications, sperm concentration, and forward progression indices. Together, these materials offer full transparency into the dataset composition, enable reproducibility of the reported results, and give additional context for interpreting the variability observed across subjects and preparation conditions.

All videos, bounding box annotations, and motility data are found at [18]: https://huggingface.co

### Calibration

Spatial calibration was performed to convert pixel-based measurements into physical units (*µ*m) for both the optical microscope and the CASA system. For each imaging system, calibration factors were extracted separately for each magnification setting by measuring a fixed physical dimension on a calibration target and mapping the corresponding pixel distances to their equivalent lengths in microns. The calibration procedures were conducted independently for the optical microscope and the CASA system and are described in detail below. All empirical calibration measurements are reported as mean pixel values with associated standard deviations, computed from automated analysis of multiple frames extracted from calibration videos. In this study, The calibration was performed independently for the optical microscope and the CASA system using the following two methods.

1. **Theoretical method:** Theoretical calibration ratios were derived from system specifications. For the optical microscope, theoretical values were computed from the camera sensor pixel pitch and the effective optical magnification. For the CASA system, theoretical calibration values were obtained directly from manufacturer-provided specifications.
2. **Empirical method:** Empirical calibration ratios were extracted from images by measuring the number of pixels of geometric features with known dimensions; we measure a dot with a known diameter to calibrate the optical microscope, and a rule-like structure containing ticks separated by a known distance to calibrate the CASA system. For each imaging system and magnification setting, calibration images were recorded and processed using a Python pipeline to extract the number of pixels of the measured structures; we report the mean pixel measurements and associated standard deviations. The number of measured pixels is then associated with a micron value to obtain the empirical micron-per-pixel calibration ratios.

Next, we describe and illustrate the theoretical and empirical calibration procedures, and validate the resulting calibration ratios using sperm cell images recorded with both the optical microscope and the CASA system.

#### Theoretical calibration method

##### Optical Microscope

The Fujifilm X–T10 camera has a pixel pitch of 4.8 *µ*m/pixel. The optical setup included a reduction lens with a magnification factor of *r* = 0.5. Under this configuration, a nominal laboratory magnification of 100× corresponded to a 20× objective lens, while a laboratory magnification of 200× corresponded to a 40× objective lens.

The spatial calibration (micron-per-pixel ratio) was computed as follows:

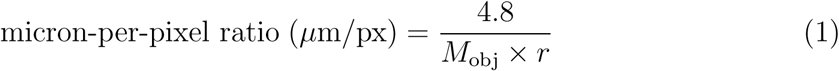

where *M*_obj_ denotes the objective magnification and *r* is the reduction factor of the optical system.

Using this formulation, the theoretical micron-per-pixel ratios were calculated as:

- 100x magnification → micron-per-pixels ratio = 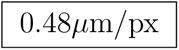
- 200x magnification → micron-per-pixels ratio = 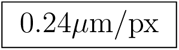

##### CASA system

The calibration ratios are directly provided by the CASA system specifications, and are:

- 200x magnification → micron-per-pixels ratio = 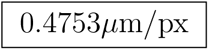
- 400x magnification → micron-per-pixels ratio = 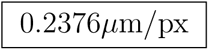

Note that the theoretical calibration ratios reported in this section were not used in the main text but are reported here for completeness. The ratios used in the main text are reported in the Empirical calibration method section.

#### Empirical calibration method

##### Optical Microscope

We used the calibration slide provided with the Swift M10DB-MP Series Advanced Microscope, shown in Figure A1. The slide contains calibration dots of known physical diameters.

**Figure A1:**
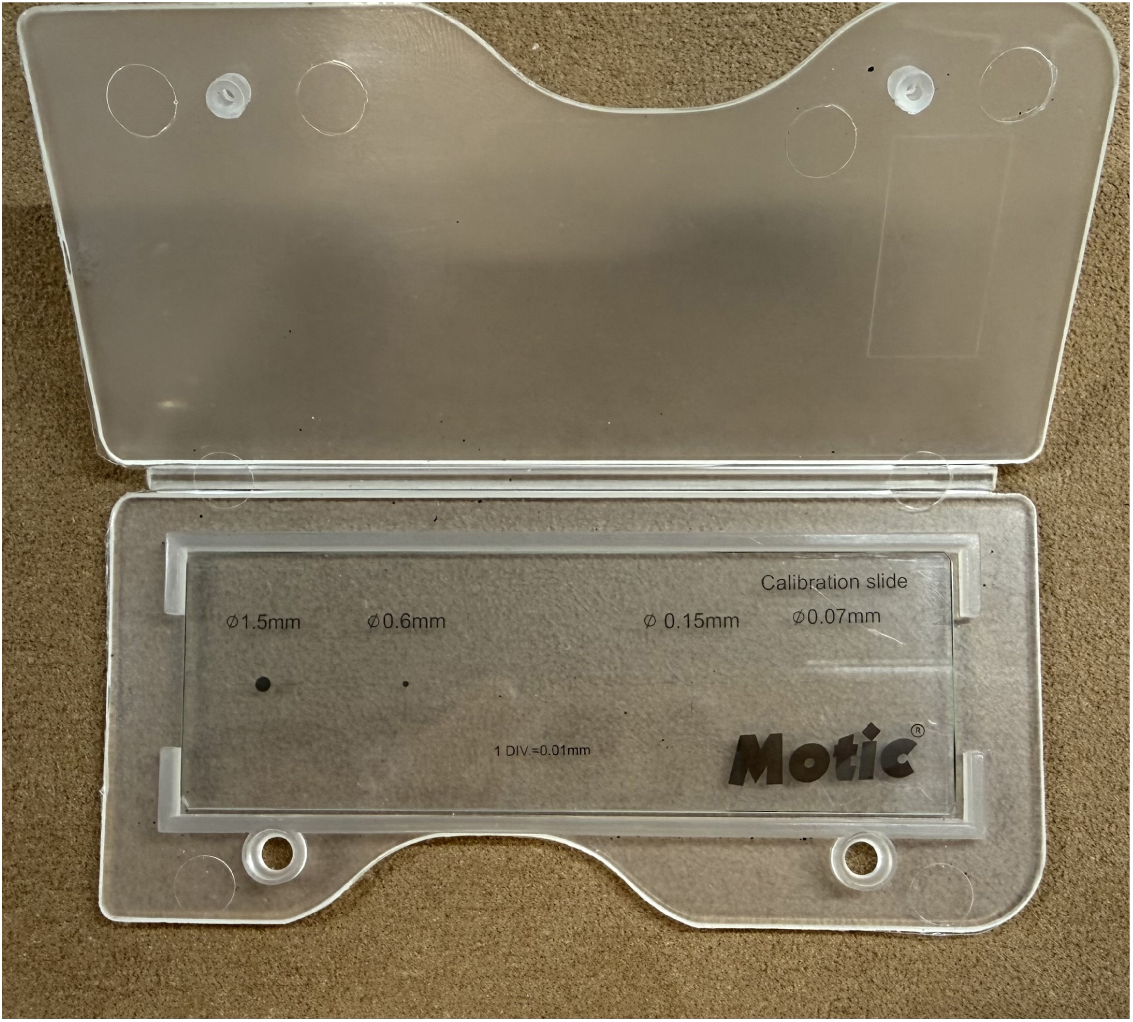
Calibration slide provided with the Swift M10DB-MP Series Advanced Microscope. The slide contains multiple calibration dots with known physical diameters (70*µ*m, 150*µ*m, 600*µ*m, and 1500*µ*m). For this application, we use the two smallest dots (70*µ*m and 150*µ*m), which are not visible to the naked eye under standard viewing conditions. These dots were used as reference targets for empirical spatial calibration of the optical microscope.

These dots were used to map pixel-based measurements of dot diameters to their corresponding physical lengths in microns. This procedure was performed independently for each magnification setting considered for the optical microscope system.

This procedure was carried out as follows:

1. Mount the calibration slide on the microscope stage.
2. Bring the 70 *µ*m reference dot into focus.
3. Acquire images using a 100× objective lens.
4. Record a video sequence of approximately 700 frames.
5. Repeat Steps 1–4 for the 150 *µ*m reference dot.
6. Repeat Steps 1–5 for the 200× objective lens.

In Table A1, we report pixel-based measurements of calibration dot diameters obtained from optical microscope images at two magnification settings (100× and 200×). For each magnification, two calibration dots of known physical diameters (70 *µ*m and 150 *µ*m) were analyzed. Table A1 lists the measured dot diameters in pixels and the corresponding predicted physical diameters in microns obtained using both theoretical and empirical calibration ratios. Theoretical calibration ratios, shown in blue, were derived from the camera pixel pitch and effective optical magnification, while empirical calibration ratios, shown in green, were estimated from image-based measurements of calibration targets with known physical dimensions.

Figure A2 shows representative calibration dot images acquired using the optical microscope at the two nominal magnification settings considered in this study (100× and 200×). For each magnification setting, calibration targets with known physical diameters of 70 *µ*m and 150 *µ*m were imaged, and calibration circles were detected to obtain pixel-based diameter measurements. For both magnification settings, pixel-based measurements obtained from the calibration targets were used to derive empirical micron-per-pixel ratios and to compare them against the corresponding theoretical calibration factors.

Based on the pixel-based calibration measurements, the following empirical micron-per-pixel ratios were adopted for use for the optical microscope system: 0.419 *µ*m/px for 100× magnification and 0.202 *µ*m/px for 200× magnification.

**Table A1:** Pixel-based measurements and predicted calibration circle diameters obtained from optical microscope images under different magnification and calibration settings. Theoretical micron-per-pixel ratios were derived from the camera pixel pitch and optical magnification model, while measured ratios were estimated empirically from image-based calibration circle measurements.

| Magnification and Calibration Setting (magnification, dot diameter) | Diameter of the calibration dot (in pixels) | Diameter of the calibration dot (in $\mu\text{m}$) | |
| --- | --- | --- | --- |
|  |  | Theoretical Ratio | Empirical Ratio |
| (a) 100×, 70 $\mu\text{m}$ | $168.67 \pm 0.14$ | $(168.67 \pm 0.14) \times 0.48$<br>$= 80.96 \pm 0.07$ | $(168.67 \pm 0.14) \times 0.415$<br>$= 69.99 \pm 0.06$ |
| (b) 100×, 150 $\mu\text{m}$ | $353.98 \pm 1.5$ | $(353.98 \pm 1.5) \times 0.48$<br>$= 169.91 \pm 0.72$ | $(353.98 \pm 1.5) \times 0.423$<br>$= 149.73 \pm 0.63$ |
| (c) 200×, 70 $\mu\text{m}$ | $350.86 \pm 0.58$ | $(350.86 \pm 0.58) \times 0.24$<br>$= 84.20 \pm 0.14$ | $(350.86 \pm 0.58) \times 0.199$<br>$= 69.82 \pm 0.12$ |
| (d) 200×, 150 $\mu\text{m}$ | $733.47 \pm 0.66$ | $(733.47 \pm 0.66) \times 0.24$<br>$= 176.03 \pm 0.16$ | $(733.47 \pm 0.66) \times 0.204$<br>$= 149.62 \pm 0.13$ |

| Case | Setting | Theoretical error ($\mu\text{m}$) | Empirical error ($\mu\text{m}$) |
| --- | --- | --- | --- |
| (a) | 100×, 70 $\mu\text{m}$ dot | $\vert 80.96 - 70\vert \pm 0.07 = 10.96 \pm 0.07$ | $\vert 69.99 - 70\vert \pm 0.06 = 0.01 \pm 0.06$ |
| (b) | 100×, 150 $\mu\text{m}$ dot | $\vert 169.91 - 150\vert \pm 0.72 = 19.91 \pm 0.72$ | $\vert 149.73 - 150\vert \pm 0.63 = 0.27 \pm 0.63$ |
| (c) | 200×, 70 $\mu\text{m}$ dot | $\vert 84.20 - 70\vert \pm 0.14 = 14.20 \pm 0.14$ | $\vert 69.82 - 70\vert \pm 0.12 = 0.18 \pm 0.12$ |
| (d) | 200×, 150 $\mu\text{m}$ dot | $\vert 176.03 - 150\vert \pm 0.16 = 26.03 \pm 0.16$ | $\vert 149.62 - 150\vert \pm 0.13 = 0.38 \pm 0.13$ |

Across all calibration settings, the empirical method yields lower absolute error than the theoretical method.

**Figure A2:**
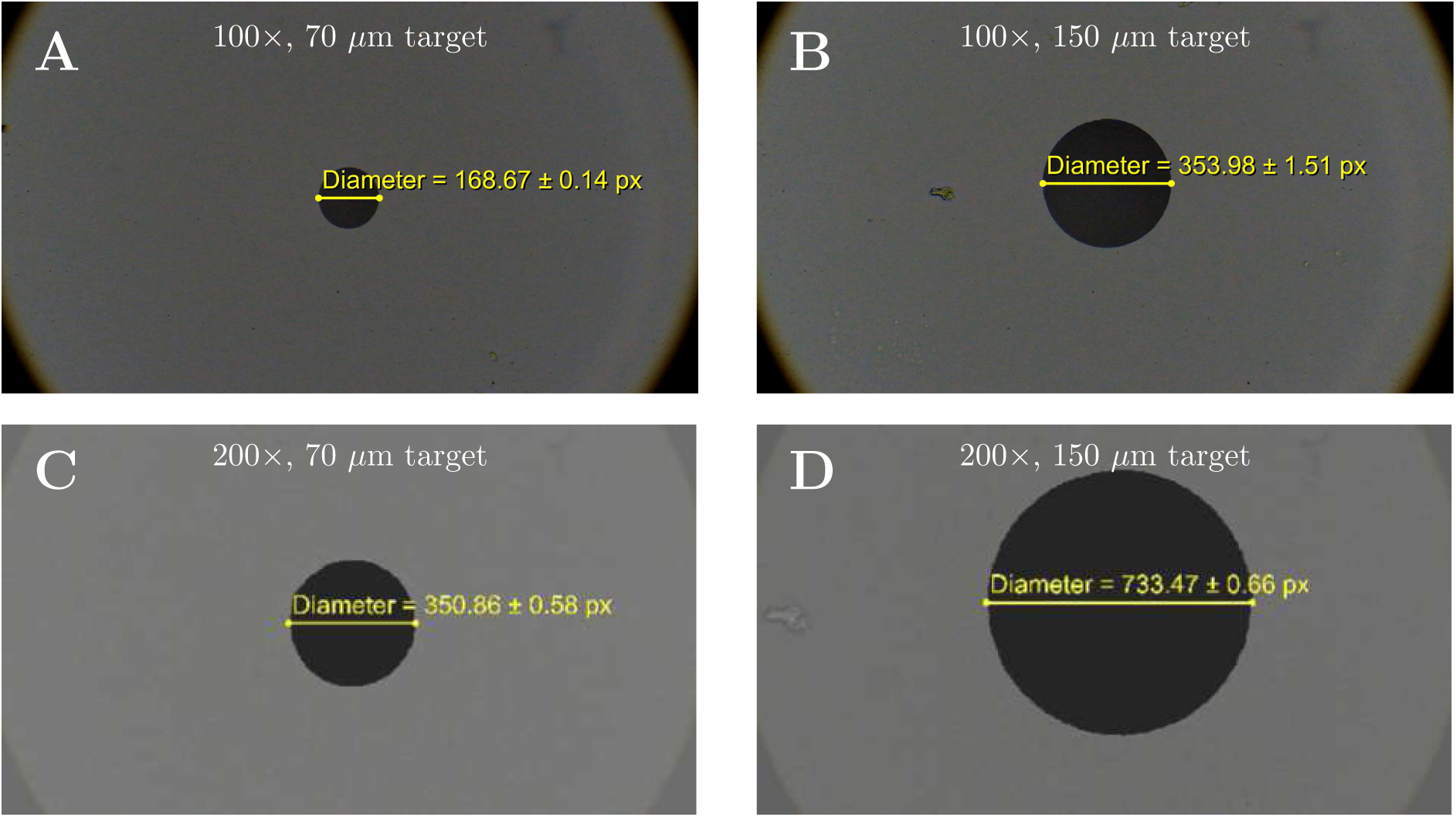
Representative calibration dot images acquired using the optical microscope at nominal magnifications of 100× and 200×. Calibration targets with known physical diameters of 70 *µ*m and 150 *µ*m are shown, with pixel-based diameter measurements used for spatial calibration.

##### CASA system

We used a Nikon stage micrometer (Model MBM11100, Type MS EA, Nikon, Japan), shown in Figure A3. The ruler contains evenly spaced calibration ticks with a known physical separation of 0.01 mm (10 microns), that are not visible to the naked eye at the acquisition scale.

These ticks were used to map pixel-based measurements of inter-tick distances to their corresponding physical lengths in microns. This procedure was performed independently for each magnification setting considered in the CASA system.

The procedure was carried out as follows.

1. Mount the calibration ruler on the CASA microscope stage.
2. Bring the calibration ruler into focus.
3. Acquire images using the 200× objective lens.
4. Record a video sequence of approximately 100 frames.
5. Measure the pixel distance between adjacent calibration ticks with a known spacing of 10 *µ*m.
6. Repeat Steps 1–5 for the 400× objective lens.

**Figure A3:**
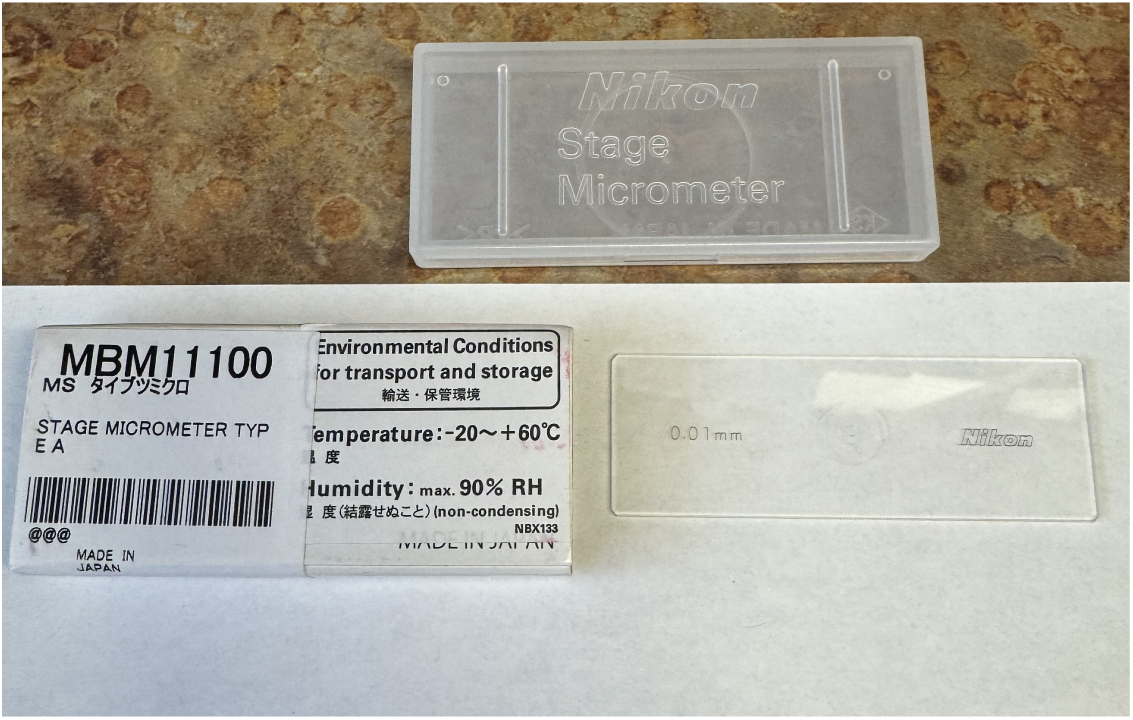
Nikon Stage Micrometer. The ruler contains evenly spaced calibration ticks with a known physical separation that are not resolvable with the naked eye. The distance between the ticks were used as reference for empirical spatial calibration of the CASA system.

In Table A2, we report pixel-based measurements of calibration tick distances obtained from CASA system images at two magnification settings (200× and 400×). For each magnification, calibration ruler ticks with a known physical spacing of 10 *µ*m were analyzed. The table lists the measured distances between adjacent calibration ticks in pixels and the corresponding predicted physical distances in microns obtained using both theoretical and empirical calibration ratios. Theoretical calibration ratios, shown in blue, were obtained from manufacturer-provided system specifications, while empirical calibration ratios, shown in green, were estimated from image-based measurements of calibration ticks with known spacing.

Figure A4 shows representative calibration ruler images acquired using the CASA system at the two nominal magnification settings considered in this study (200× and 400×). Calibration ruler ticks with a known physical spacing of 10 *µ*m were imaged, and the distances between adjacent ticks were measured to obtain pixel-based spacing values. These measurements were used to derive empirical micron-per-pixel ratios and to compare them against the corresponding theoretical calibration factors.

**Table A2:** Pixel-based measurements and predicted calibration tick distances obtained from the CASA system under different magnification settings. Theoretical micron-per-pixel ratios were derived from system specifications, while empirical ratios were estimated from image-based calibration tick measurements. Uncertainties in micron-scale distances were obtained by linear propagation of pixel-level measurement uncertainty.

| Magnification and Calibration Setting<br>(magnification, tick spacing) | Distance between calibration ticks<br>(in pixels) | Distance between calibration ticks (in $\mu\text{m}$) | |
| --- | --- | --- | --- |
|  |  | Theoretical Ratio | Empirical Ratio |
| 200×, 10 $\mu\text{m}$ | $25.187 \pm 0.12$ | $(25.187 \pm 0.12) \times 0.4753$<br>$= 11.97 \pm 0.06$ | $(25.187 \pm 0.12) \times 0.3970$<br>$= 9.99 \pm 0.05$ |
| 400×, 10 $\mu\text{m}$ | $50.571 \pm 0.60$ | $(50.571 \pm 0.60) \times 0.2376$<br>$= 12.02 \pm 0.14$ | $(50.571 \pm 0.60) \times 0.1977$<br>$= 10.00 \pm 0.12$ |

| Case | Setting | Theoretical error ($\mu\text{m}$) | Empirical error ($\mu\text{m}$) |
| --- | --- | --- | --- |
| (a) | 200×, 10 $\mu\text{m}$ tick spacing | $\vert 11.97 - 10\vert \pm 0.06 = 1.97 \pm 0.06$ | $\vert 9.99 - 10\vert \pm 0.05 = 0.01 \pm 0.05$ |
| (b) | 400×, 10 $\mu\text{m}$ tick spacing | $\vert 12.02 - 10\vert \pm 0.14 = 2.02 \pm 0.14$ | $\vert 10.00 - 10\vert \pm 0.12 = 0.00 \pm 0.12$ |

Across both magnification settings, the empirical method yields lower absolute error than the theoretical method.

**Figure A4:**
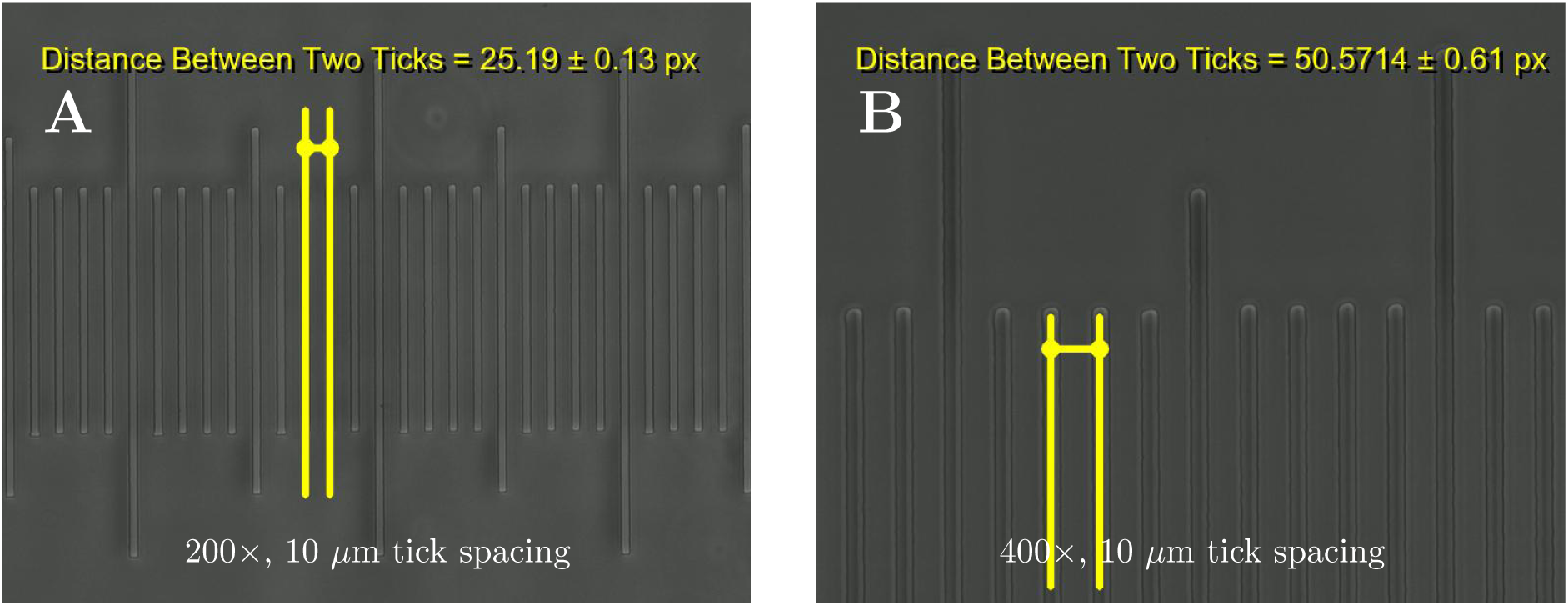
Representative calibration ruler images acquired using the CASA system at nominal magnifications of 200× and 400×. Calibration ruler ticks with a known physical spacing of 10 *µ*m are shown, with pixel-based inter-tick distance measurements used to derive empirical micron-per-pixel calibration ratios. Annotated overlays indicate representative tick pairs used for visualization.

Based on the pixel-based calibration measurements, the following empirical micron-per-pixel ratios were used for the CASA system: 0.397 *µ*m/px for 200× magnification and 0.198 *µ*m/px for 400× magnification.

##### Example Images

To illustrate the practical application of the spatial calibration procedures described in the Empirical calibration method section, this section presents representative example images. They demonstrate how pixel-based measurements are converted into physical dimensions in sperm head analysis.

Figures A5 and A6 show representative example images acquired using the optical microscope at 100× and 200× nominal magnifications, respectively. In each image, a detected sperm head is annotated with its major and minor axes, and the corresponding physical dimensions are computed using both the theoretical micron-per-pixel ratios derived from the optical calibration model and the empirically measured ratios obtained from calibration targets. These examples illustrate how the optical microscope calibration factors are applied to convert pixel-level measurements into physical units under different magnification settings.

Figures A7 and A8 show corresponding example images acquired using the CASA system at 200× and 400× nominal magnifications, illustrating the application of CASA-specific calibration ratios to the same downstream morphometric measurements.

**Figure A5:**
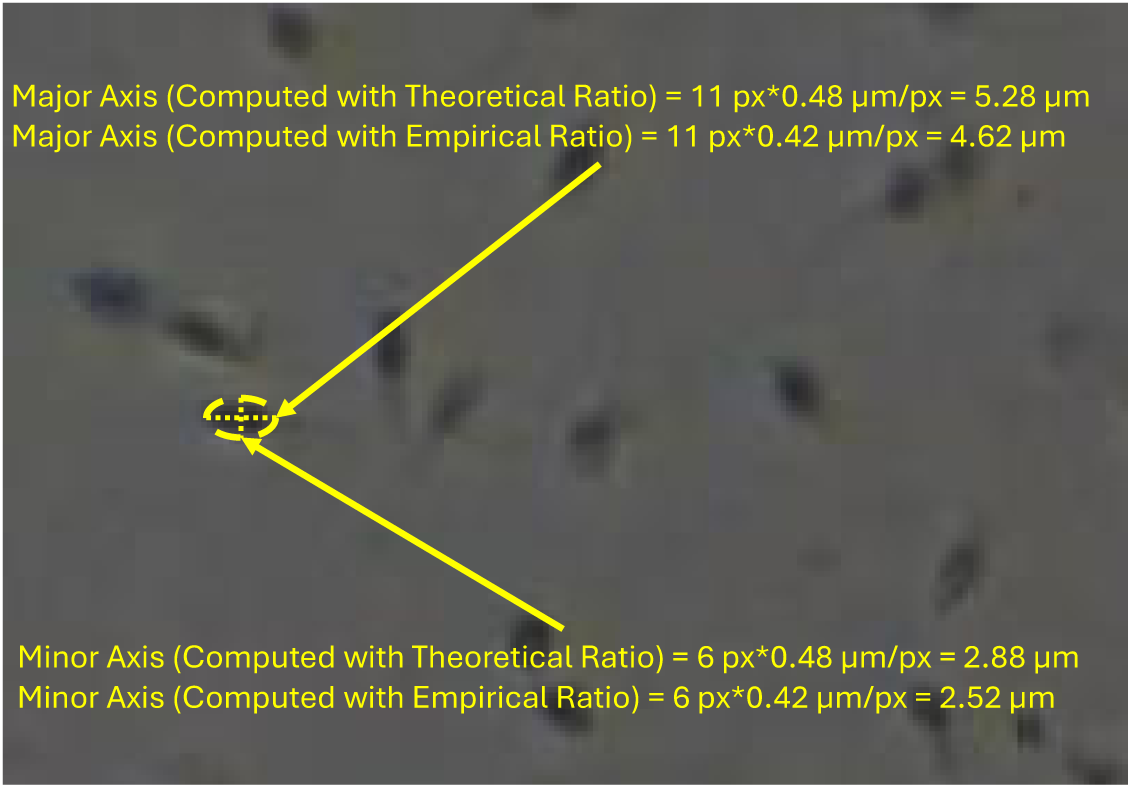
Representative optical microscope image acquired at 100× nominal magnification. A detected sperm head is annotated with its major and minor axes, and physical dimensions are computed using both the theoretical micron-per-pixel ratio from the optical calibration model and the empirically measured ratio.

**Figure A6:**
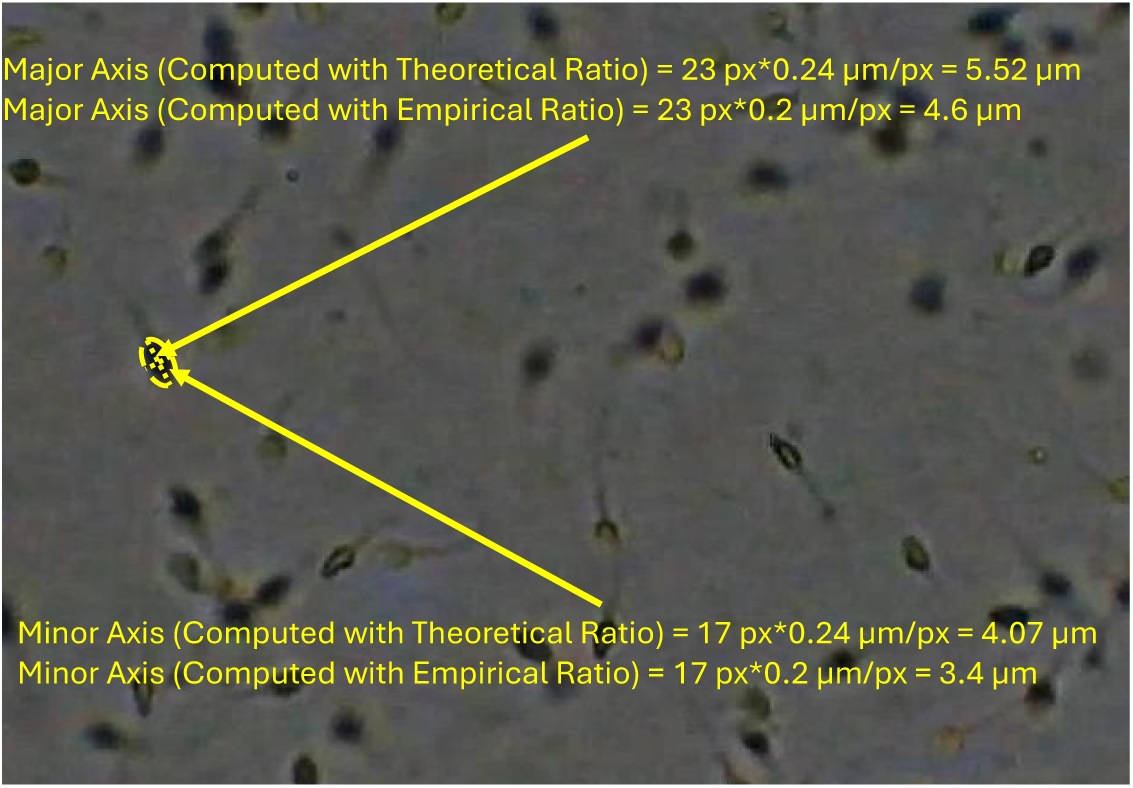
Representative optical microscope image acquired at 200× nominal magnification. The detected sperm head is annotated with major and minor axis lengths computed using theoretical and empirically measured micron-per-pixel calibration ratios.

**Figure A7:**
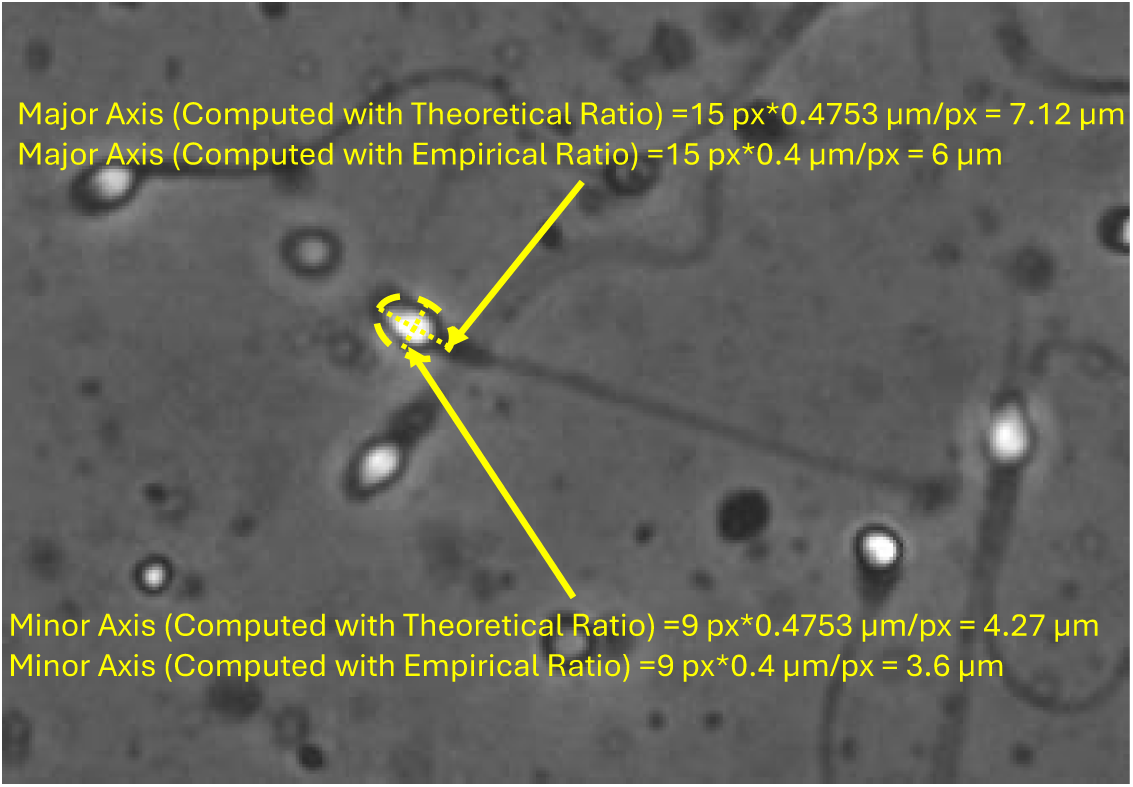
Representative CASA system image acquired at 200× nominal magnification. Major and minor axes of a detected sperm head are annotated, with physical dimensions computed using manufacturer-specified (theoretical) and empirically measured micron-per-pixel ratios.

**Figure A8:**
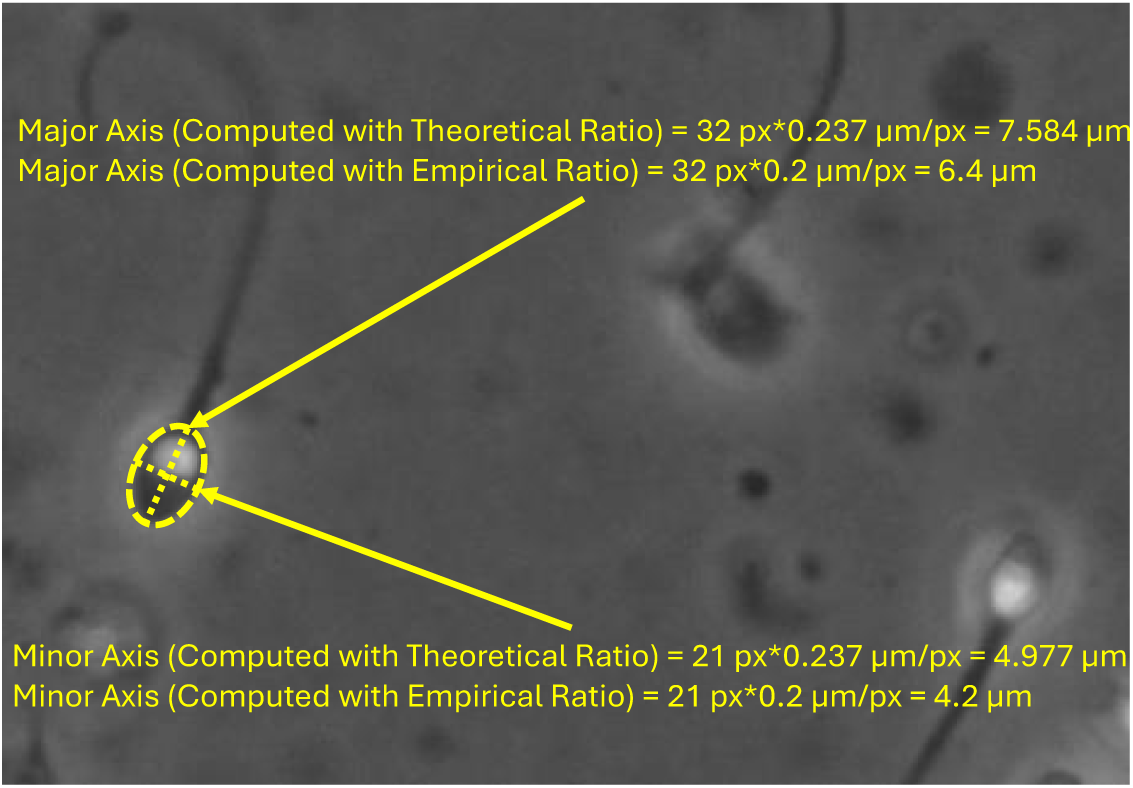
Representative CASA system image acquired at 400× nominal magnification. The detected sperm head is annotated with major and minor axes, and physical dimensions are computed using theoretical calibration values from system specifications and empirically measured calibration ratios obtained from pixel-based analysis.

Overall, the calibrated physical dimensions obtained from these examples are consistent with nominal sperm head size ranges reported in the literature, supporting the plausibility of the adopted micron-per-pixel calibration values.

### Detection and Tracking Experiments

This appendix reports in full the detection, tracking, and kinematic-parameter experiments summarized in the Technical Validation section. Sperm heads were detected with YOLOv5, associated across frames with SORT to form trajectories, and the trajectories were then used to compute the eight kinematic parameters defined in the Motility Parameters section. The same pipeline was applied to the manually annotated ground truth, so that parameters derived from automatic detections and from the annotations can be compared directly. Together these experiments illustrate the range of tasks the dataset supports.

#### Object Detection using YOLOv5

We applied the YOLOv5 object detection framework [14] to both the CASA and optical microscope recordings. YOLOv5 is a state-of-the-art object detector capable of fast and accurate detections. Its efficiency and speed make it well-suited for frame-by-frame sperm cell localization.

For the CASA recordings, we used the pretrained YOLOv5m weights released as part of the VISEM-Tracking project [20]. The VISEM-Tracking model was originally trained on approximately 685,000 annotated sperm cells from 29,196 frames, split 80/20 into a training set (525,000 annotated sperm cells, 23,357 frames) and a testing set (131,000 annotated sperm cells, 5,839 frames).

In this context, training refers to the iterative optimization of model parameters using annotated data to minimize detection errors and improve generalization to unseen samples. The training involved image resizing to 640×640 pixels, batch size 16, and 300 epochs, with all other hyper parameters at YOLOv5 defaults. Training was performed on two NVIDIA GeForce RTX 3080 GPUs with an AMD Ryzen 9 3950X CPU. Table A3 displays the performance metrics for the YOLOv5 models reported on the VISEM-Tracking dataset [20]. Applying these weights to our manually annotated CASA subset shows how a model trained on a different dataset performs on these recordings.

For the optical microscope recordings, we trained different-sized YOLOv5 models from scratch on our manually annotated subset, which contains approximately 600,000 annotated sperm cells from 17,300 frames. The data were split 80/20 into a training set (16 videos, 396,262 annotated sperm cells, 13,743 frames) and a testing set (4 videos, 209,611 annotated sperm cells, 3,607 frames). The training involved image resizing in to 832x832 pixels, batch size 16, and used 100 epochs, with other hyperparameters left at YOLOv5 defaults. Training was performed on an Intel Core i9-10900K CPU, 32 GB RAM, and an NVIDIA RTX Quadro 4000 GPU. Table A3 summarizes the performance of the five YOLOv5 variants trained on our optical microscope dataset. As shown in the table, precision values range from 0.64 (YOLOv5s) to 0.66 (YOLOv5n), while recall is consistently higher across models (0.72-0.76). The smallest (YOLOv5n) achieved the highest mAP0.5:0.95 score (0.41) and overall fitness value (0.37).

**Table A3:**
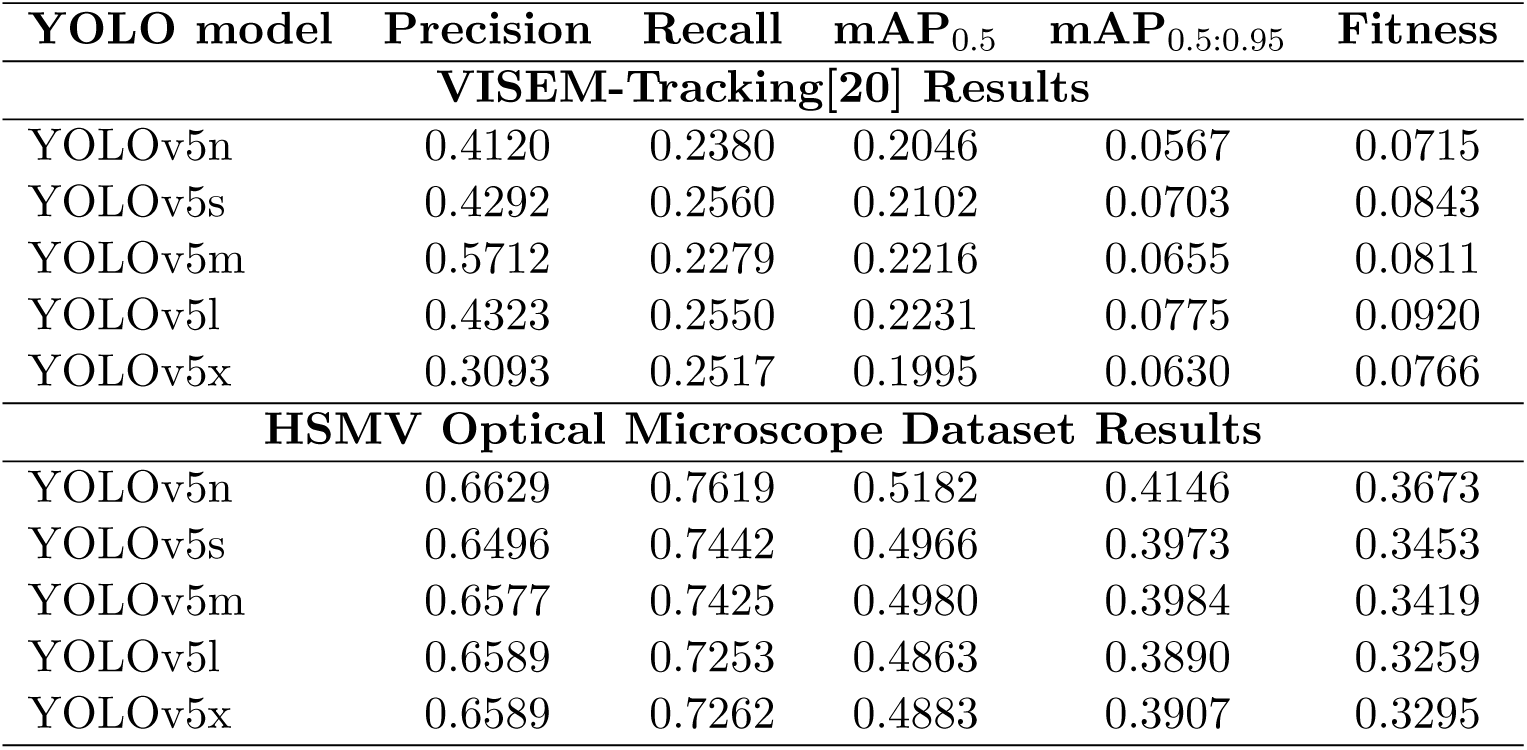
Comparison of YOLOv5 performance metrics between the VISEM-Tracking[20] models and the models trained on our optical microscope dataset.

| **YOLO model** | **Precision** | **Recall** | **mAP<sub>0.5</sub>** | **mAP<sub>0.5:0.95</sub>** | **Fitness** |
| --- | --- | --- | --- | --- | --- |
| **VISEM-Tracking[20] Results** |  |  |  |  |  |
| YOLOv5n | 0.4120 | 0.2380 | 0.2046 | 0.0567 | 0.0715 |
| YOLOv5s | 0.4292 | 0.2560 | 0.2102 | 0.0703 | 0.0843 |
| YOLOv5m | 0.5712 | 0.2279 | 0.2216 | 0.0655 | 0.0811 |
| YOLOv5l | 0.4323 | 0.2550 | 0.2231 | 0.0775 | 0.0920 |
| YOLOv5x | 0.3093 | 0.2517 | 0.1995 | 0.0630 | 0.0766 |
| **HSMV Optical Microscope Dataset Results** |  |  |  |  |  |
| YOLOv5n | 0.6629 | 0.7619 | 0.5182 | 0.4146 | 0.3673 |
| YOLOv5s | 0.6496 | 0.7442 | 0.4966 | 0.3973 | 0.3453 |
| YOLOv5m | 0.6577 | 0.7425 | 0.4980 | 0.3984 | 0.3419 |
| YOLOv5l | 0.6589 | 0.7253 | 0.4863 | 0.3890 | 0.3259 |
| YOLOv5x | 0.6589 | 0.7262 | 0.4883 | 0.3907 | 0.3295 |

The YOLOv5 model trained on optical microscope recordings achieved a precision of 0.66 and a recall of 0.76 on the held-out 20% testing set. Applying the same YOLOv5 weights trained on the VISEM-Tracking dataset to our testing CASA subset resulted precision of 0.82 and a recall of 0.43, likely reflecting differences in dataset characteristics.

Overall, these results demonstrate that our dataset enables effective training of YOLOv5 models for sperm detection. Although stricter localization metrics (mAP_0.5_, mAP_0.5:0.95_, and fitness) reflect the challenges inherent to microscopic imagery, the detected bounding boxes are sufficiently precise for subsequent analysis. In the Motility Parameters section, we show that the detections produced by YOLOv5 support accurate motility parameter estimation within a complete analysis pipeline.

#### Tracking using Simple Online and Realtime Tracking (SORT)

We employed the Simple Online and Realtime Tracking (SORT) algorithm [4] to generate sperm cell trajectories from both YOLOv5 detections and ground truth bounding boxes. SORT is a lightweight tracking algorithm that combines a Kalman filter for motion prediction with the Hungarian algorithm for data association, using Intersection-over-Union (IoU) [15] as the cost metric.

In our workflow, SORT was executed in two modes. In the first mode, ground truth bounding boxes were used as inputs to produce reference trajectories. In the second mode, YOLOv5 detections were used as inputs, enabling us to benchmark the accuracy of tracking pipelines that rely on automatic detections. This dual-mode approach allows for a direct visual comparison between automated tracking results and manually verified ground truth.

Figures A9 and A10 show the detections and tracks obtained from frames 10, 20, 30, and 40 of participant sys-casa sub-HC011 ses-01 run-003 for the CASA recordings and frames 10, 20, 30, and 40 of participant sys-opt sub-HC003 ses-01 run-031 for the optical microscope recordings, respectively. Trajectories derived from ground-truth annotations are shown in red, while those based on YOLOv5 detections are shown in cyan. These examples illustrate the capability of YOLOv5, followed by the SORT pipelines, to approximate ground truth trajectories, exemplifying the utility of the HSMV dataset for developing and evaluating sperm tracking methods.

#### Motility Parameters

We computed sperm motility parameters from the tracks calculated using SORT from both YOLOv5-based detections and manually annotated ground truth. These parameters are widely used in clinical and research settings to characterize sperm movements.

**Figure A9:**
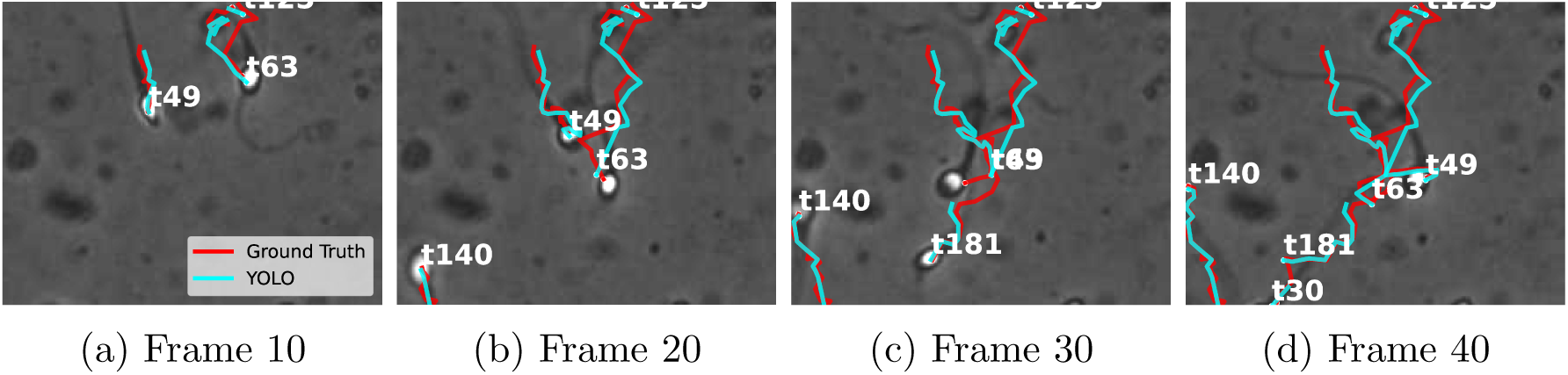
Detections and tracks obtained from the CASA recordings (participant sys-casa sub-HC011 ses-01 run-003). Ground truth-based trajectories (red) vs YOLOv5-based trajectories (cyan) at frames 10, 20, 30, and 40.

**Figure A10:**
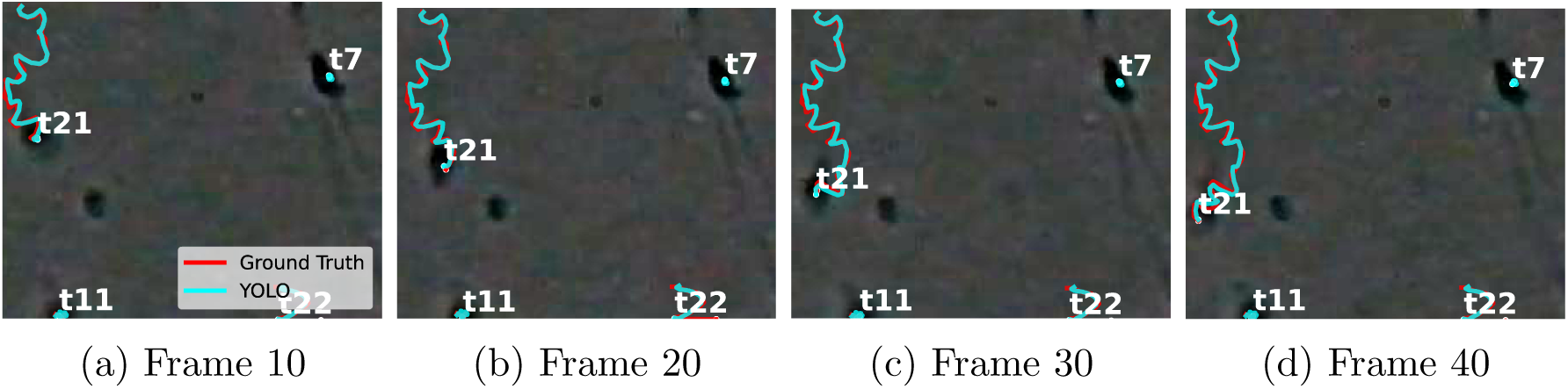
Detections and tracks obtained from the optical microscope recordings (participant sys-opt sub-HC003 ses-01 run-031). Ground truth-based trajectories (red) vs YOLOv5-based trajectories (cyan) at frames 10, 20, 30, and 40.

We extracted the following eight standard motility parameters from sperm trajectories as was done in [21]:

- Curvilinear Velocity (VCL): The total distance traveled by a sperm cell divided by the time duration.
- Straight-Line Velocity (VSL): The straight-line distance between the first and last detected positions divided by time.
- Average Path Velocity (VAP): The average velocity along a smoothed trajectory.
- Linearity (LIN): Defined as VSL divided by VCL, indicating how straight the path is.
- Wobble (WOB): Defined as VAP divided by VCL, measuring oscillations around the average path.
- Straightness (STR): Defined as VSL divided by VAP, indicating the straightness of the smoothed path.
- Amplitude of Lateral Head Displacement (ALH): The average deviation of a sperm cell from its average path.
- Mean Angular Displacement (MAD): The average turning angle of the sperm cell along its curvilinear path.

We computed density scatter plots of pairwise motility parameters by placing two features on the horizontal and vertical axes and encoding point density with a color gradient (warmer colors indicate higher densities). Figures A11 and A12 show the motility parameters computed from ground truth and YOLOv5-based trajectories, respectively, for subject sys-casa sub-HC006 ses-01 run-005 of the CASA system recordings. Additionally, Figures A13 and A14 show the motility parameters computed from ground truth and YOLOv5-based trajectories, respectively, for subject sys-opt sub-HC003 ses-01 run-025 of the optical microscope recordings.

For the CASA recordings, the agreement between the YOLOv5-derived and ground truth motility patterns was strong across all evaluated parameter pairs. Using Pearson correlations computed between the corresponding two-dimensional density histograms, the VCL-VSL, LIN-ALH, WOB-VSL, and LIN-MAD pairs yielded correlation values ranging from 0.83 to 0.89 (all p *<* 0.001), indicating that YOLOv5 closely reproduces the ground truth based motility distributions. The mean values of the corresponding parameters were also similar between the motility parameters computed from the ground truth trajectories and YOLOv5, further confirming consistency between the two methods. For the optical microscope recordings, the LIN-ALH, WOB-VSL, and LIN-MAD pairs showed moderate-to-strong correspondence (r = 0.64-0.67, all p *<* 0.001) with closely aligned mean values. Although the VCL-VSL pair exhibited lower density correlation (r = 0.30, p = 0.001), the mean values of the motility parameters remained highly comparable between the ground truth and YOLOv5 estimates, indicating that the measurements remain practically consistent and usable despite the lower distribution-level agreement relative to the CASA recordings.

To complement the density scatter plots, Figure 3 shows spider plots summarizing the average values of the eight motility parameters across four representative participants from the CASA and optical microscope systems. In the comparisons, the close overlap YOLOv5 (orange) and ground truth (blue) plots illustrates the behavior of the automated pipeline.

**Figure A11:**
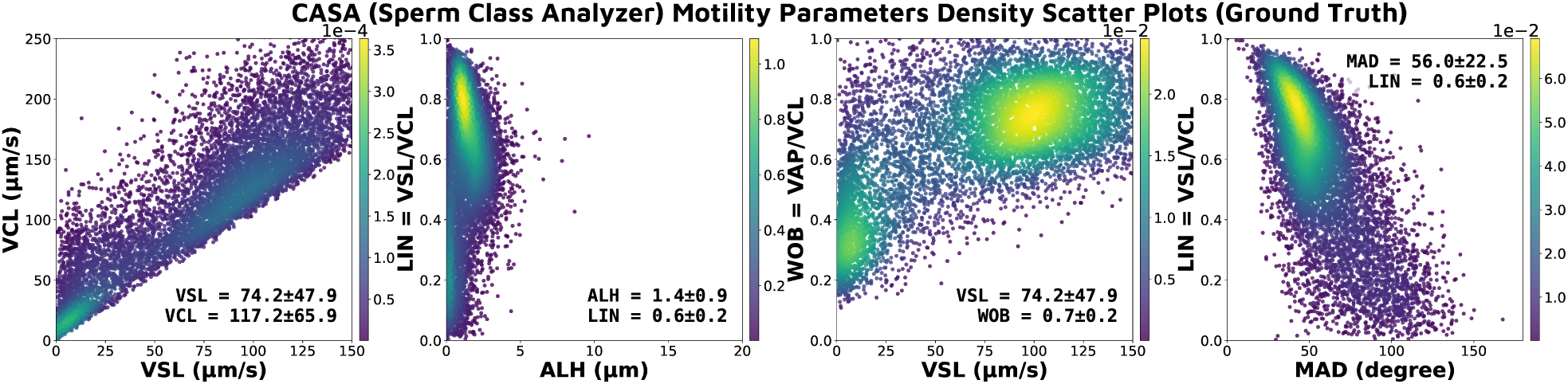
CASA recordings (participant sys-casa sub-HC006) motility parameters visualized as density scatter plots using ground truth trajectories. Each panel shows pairwise relationships between motility parameters.

**Figure A12:**
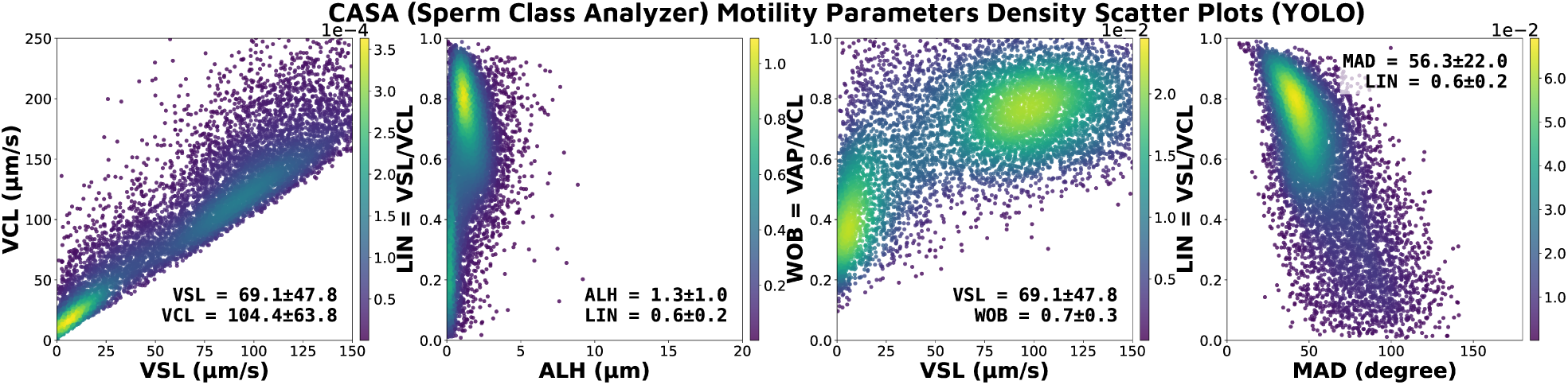
CASA recordings (participant sys-casa sub-HC006) motility parameters visualized as density scatter plots using YOLOv5 detections. Each panel shows pairwise relationships between motility parameters.

### Dataset Property Tables

**Figure A13:**
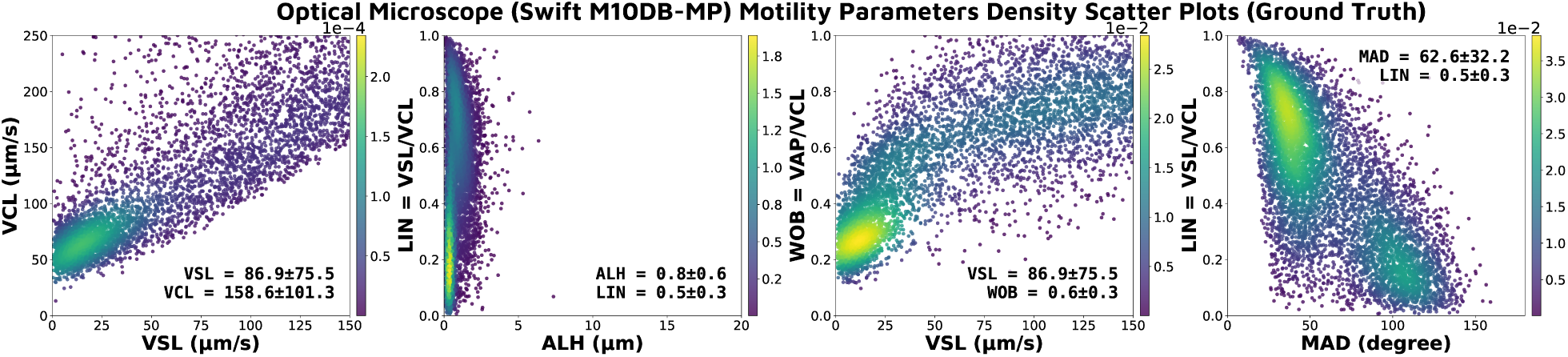
Optical microscope recordings (participant sys-opt sub-HC003) motility parameters visualized as density scatter plots using ground truth trajectories. Each panel shows pairwise relationships between motility parameters.

**Figure A14:**
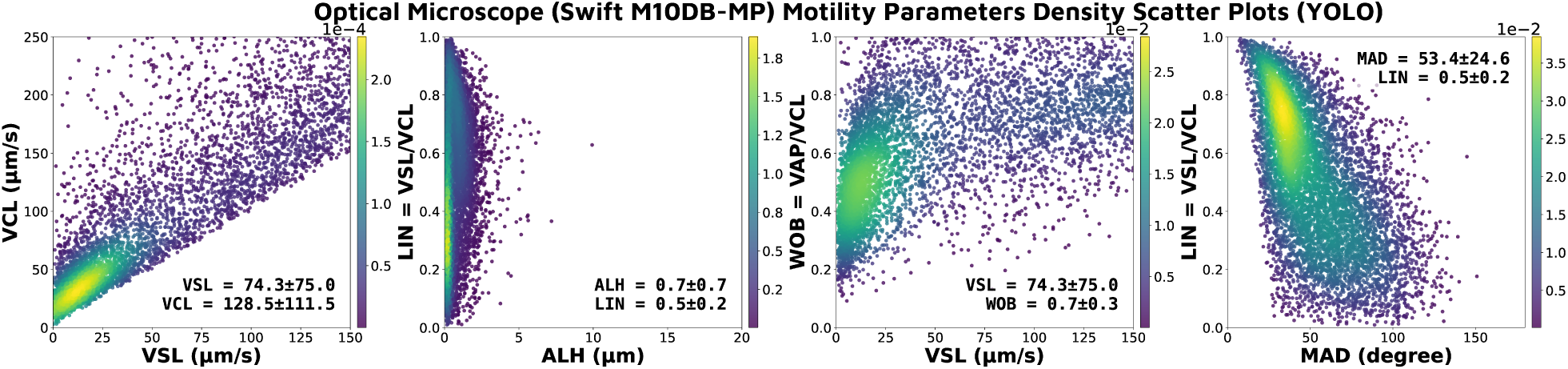
Optical microscope recordings (participant sys-opt sub-HC003) motility parameters visualized as density scatter plots using YOLOv5 detections. Each panel shows pairwise relationships between motility parameters.

**Table A4:**
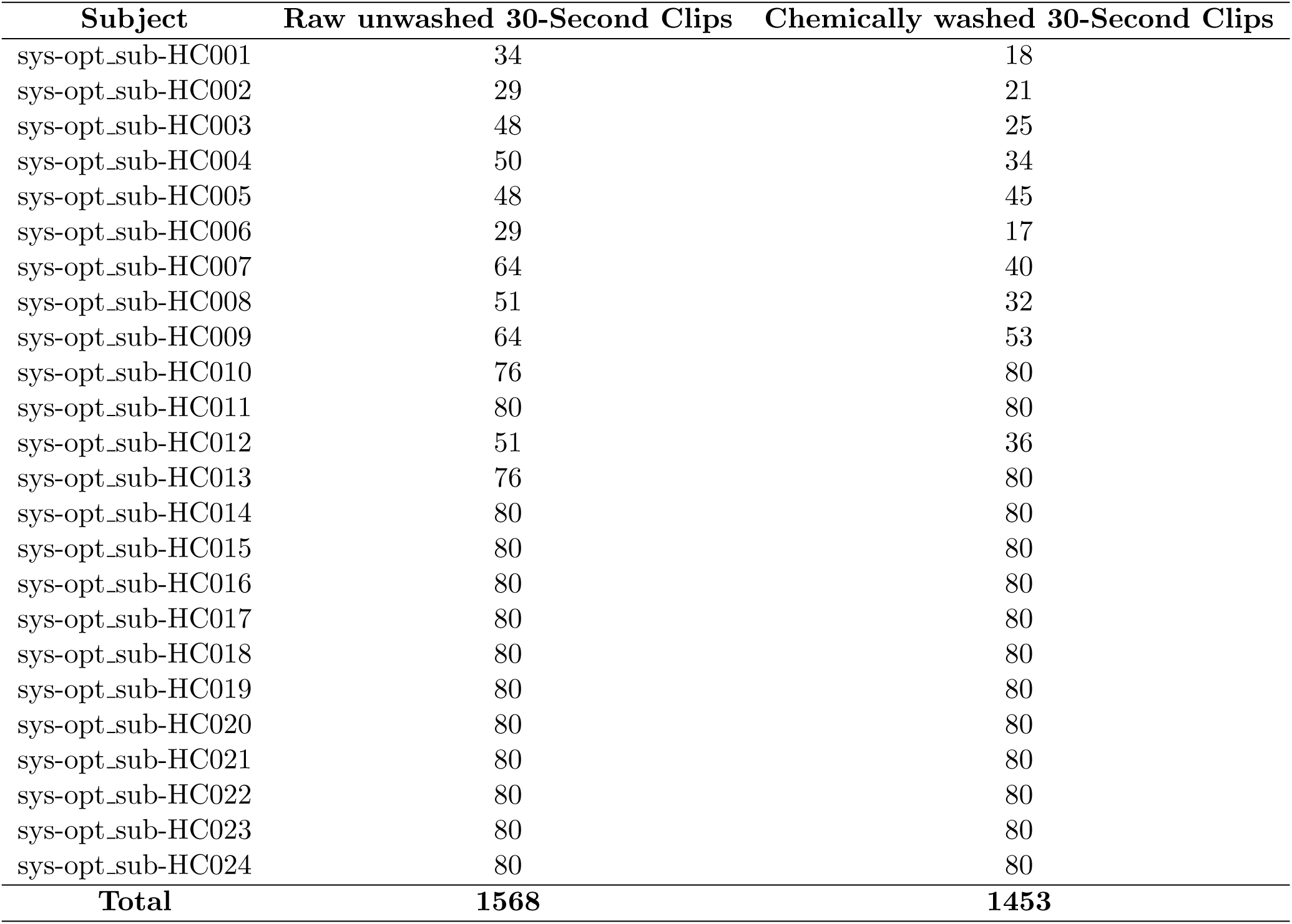
Optical microscope video clips

**Table A5:** CASA video clips

| Subject | Raw unwashed 30-Second Clips | Chemically washed 30-Second Clips |
| --- | --- | --- |
| sys-casa_sub-HC001 | 2 | 2 |
| sys-casa_sub-HC002 | 2 | 2 |
| sys-casa_sub-HC003 | 4 | 4 |
| sys-casa_sub-HC004 | 2 | 4 |
| sys-casa_sub-HC005 | 2 | 2 |
| sys-casa_sub-HC006 | 4 | 4 |
| sys-casa_sub-HC007 | 2 | 2 |
| sys-casa_sub-HC008 | 4 | 4 |
| sys-casa_sub-HC009 | 2 | 2 |
| sys-casa_sub-HC010 | 2 | 2 |
| sys-casa_sub-HC011 | 2 | 2 |
| sys-casa_sub-HC012 | 2 | 2 |
| sys-casa_sub-HC013 | 2 | 2 |
| sys-casa_sub-HC014 | 2 | 2 |
| sys-casa_sub-HC015 | 2 | 2 |
| sys-casa_sub-HC016 | 2 | 4 |
| sys-casa_sub-HC017 | 2 | 4 |
| sys-casa_sub-HC018 | 2 | 4 |
| sys-casa_sub-HC019 | 2 | 2 |
| sys-casa_sub-HC020 | 2 | 4 |
| sys-casa_sub-HC021 | 2 | 3 |
| sys-casa_sub-HC022 | 3 | 3 |
| sys-casa_sub-HC023 | 3 | 2 |
| sys-casa_sub-HC024 | 3 | 2 |
| sys-casa_sub-HC025 | 3 | 2 |
| sys-casa_sub-HC026 | 2 | 2 |
| sys-casa_sub-HC027 | 3 | 5 |
| **Total** | **65** | **73** |

**Table A6:** Summary of total sperm cells and total annotated frames for 14 CASA subjects

| Subject | Total Annotated Sperm Cells | Total Annotated Frames |
| --- | --- | --- |
| sys-casa_sub-HC004 | 81179 | 900 |
| sys-casa_sub-HC006 | 39652 | 900 |
| sys-casa_sub-HC008 | 8318 | 900 |
| sys-casa_sub-HC009 | 29757 | 900 |
| sys-casa_sub-HC010 | 68386 | 900 |
| sys-casa_sub-HC011 | 14073 | 900 |
| sys-casa_sub-HC012 | 10513 | 900 |
| sys-casa_sub-HC013 | 7388 | 900 |
| sys-casa_sub-HC014 | 30699 | 900 |
| sys-casa_sub-HC015 | 27897 | 900 |
| sys-casa_sub-HC016 | 119163 | 900 |
| sys-casa_sub-HC018 | 100122 | 900 |
| sys-casa_sub-HC021 | 55059 | 900 |
| sys-casa_sub-HC027 | 95992 | 900 |
| **Total** | **688198** | **12600** |

**Table A7:** Summary of total sperm cells and total annotated frames for 20 optical microscope subjects

| **Subject** | **Total Annotated Sperm Cells** | **Total Annotated Frames** |
| --- | --- | --- |
| sys-opt_sub-HC001 | 13216 | 900 |
| sys-opt_sub-HC002 | 51843 | 200 |
| sys-opt_sub-HC003 | 19126 | 900 |
| sys-opt_sub-HC004 | 41850 | 900 |
| sys-opt_sub-HC005 | 20832 | 900 |
| sys-opt_sub-HC006 | 78357 | 934 |
| sys-opt_sub-HC007 | 4500 | 900 |
| sys-opt_sub-HC008 | 99273 | 900 |
| sys-opt_sub-HC009 | 12394 | 900 |
| sys-opt_sub-HC010 | 132263 | 900 |
| sys-opt_sub-HC011 | 15888 | 900 |
| sys-opt_sub-HC012 | 21091 | 900 |
| sys-opt_sub-HC013 | 23656 | 900 |
| sys-opt_sub-HC014 | 63147 | 900 |
| sys-opt_sub-HC015 | 61264 | 900 |
| sys-opt_sub-HC016 | 6572 | 900 |
| sys-opt_sub-HC017 | 26450 | 900 |
| sys-opt_sub-HC018 | 76235 | 900 |
| sys-opt_sub-HC019 | 18724 | 900 |
| sys-opt_sub-HC020 | 15996 | 916 |
| **Total** | **802677** | **17356** |

**Table A8:** CASA Pre-Prep Motility Parameters

| Subject | Collection Date | Vol (mL) | % Rapid | % Slow | % Non-prog | % Immotile | Sperm Conc. ($10^6$/mL) |
| --- | --- | --- | --- | --- | --- | --- | --- |
| sys-casa_sub-HC001 | 2022-01-13 | 2.0 | 37 | 22 | 13 | 28 | 55 |
| sys-casa_sub-HC002 | 2022-01-13 | 8.0 | 25 | 15 | 8 | 52 | 40 |
| sys-casa_sub-HC003 | 2022-01-18 | 3.2 | 16 | 25 | 7 | 52 | 214 |
| sys-casa_sub-HC004 | 2022-01-18 | 2.2 | 42 | 23 | 9 | 26 | 69 |
| sys-casa_sub-HC005 | 2022-01-20 | 4.5 | 65 | 15 | 1 | 19 | 8 |
| sys-casa_sub-HC006 | 2022-01-20 | 1 | 45 | 13 | 5 | 37 | 148 |
| sys-casa_sub-HC007 | 2022-01-24 | 6.1 | 33 | 32 | 5 | 30 | 22 |
| sys-casa_sub-HC008 | 2022-01-24 | 2.5 | 47 | 10 | 4 | 39 | 75 |
| sys-casa_sub-HC009 | 2022-01-24 | 3.6 | 65 | 11 | 5 | 19 | 28 |
| sys-casa_sub-HC010 | 2022-01-26 | 2.3 | 79 | 7 | 5 | 9 | 40 |
| sys-casa_sub-HC011 | 2022-01-26 | 1.1 | 7 | 21 | 8 | 64 | 48 |
| sys-casa_sub-HC012 | 2022-01-26 | 4.5 | 40 | 4 | 4 | 52 | 7 |
| sys-casa_sub-HC013 | 2022-02-01 | 3 | 10 | 22 | 10 | 58 | 5 |
| sys-casa_sub-HC014 | 2022-02-01 | 3.3 | 26 | 23 | 8 | 43 | 32 |
| sys-casa_sub-HC015 | 2022-02-01 | 3.5 | 26 | 22 | 8 | 44 | 30 |
| sys-casa_sub-HC016 | 2022-02-25 | 3 | 44 | 18 | 10 | 28 | 190 |
| sys-casa_sub-HC017 | 2022-02-25 | 4.5 | 60 | 20 | 4 | 14 | 110 |
| sys-casa_sub-HC018 | 2022-02-25 | 2.2 | 61 | 10 | 8 | 21 | 348 |
| sys-casa_sub-HC019 | 2022-03-04 | 3.6 | 35 | 25 | 10 | 30 | 57 |
| sys-casa_sub-HC020 | 2022-03-04 | 1 | 38 | 8 | 12 | 42 | 132 |
| sys-casa_sub-HC021 | 2022-03-09 | 3.3 | 46 | 23 | 3 | 28 | 52 |
| sys-casa_sub-HC022 | 2022-03-09 | 2 | 28 | 15 | 5 | 52 | 633 |
| sys-casa_sub-HC023 | 2022-03-09 | 1.5 | 12 | 18 | 12 | 58 | 86 |
| sys-casa_sub-HC024 | 2022-03-11 | 2 | 54 | 6 | 5 | 35 | 18 |
| sys-casa_sub-HC025 | 2022-03-18 | 1.5 | 33 | 36 | 8 | 23 | 26 |
| sys-casa_sub-HC026 | 2022-03-18 | 3 | 28 | 24 | 6 | 42 | 14 |
| sys-casa_sub-HC027 | 2022-03-18 | 3.1 | 63 | 7 | 4 | 26 | 64 |

**Table A9:** CASA Post-Prep Motility Parameters

| Subject | Collection Date | Vol (mL) | % Rapid | % Slow | % Non-prog | % Immotile | Sperm Conc. (10<sup>6</sup>/mL) |
| --- | --- | --- | --- | --- | --- | --- | --- |
| sys-casa_sub-HC001 | 2022-01-13 | 0.5 | 39 | 27 | 11 | 23 | 84 |
| sys-casa_sub-HC002 | 2022-01-13 | 0.5 | 55 | 25 | 8 | 12 | 128 |
| sys-casa_sub-HC003 | 2022-01-18 | 0.5 | 34 | 24 | 9 | 33 | 175 |
| sys-casa_sub-HC004 | 2022-01-18 | 0.5 | 57 | 28 | 6 | 9 | 188 |
| sys-casa_sub-HC005 | 2022-01-20 | 0.5 | 88 | 9 | 2 | 1 | 87 |
| sys-casa_sub-HC006 | 2022-01-20 | 0.5 | 80 | 10 | 3 | 7 | 288 |
| sys-casa_sub-HC007 | 2022-01-24 | 0.5 | 41 | 18 | 5 | 36 | 43 |
| sys-casa_sub-HC008 | 2022-01-24 | 0.5 | 80 | 14 | 3 | 3 | 85 |
| sys-casa_sub-HC009 | 2022-01-24 | 0.5 | 70 | 14 | 4 | 12 | 22 |
| sys-casa_sub-HC010 | 2022-01-26 | 0.5 | 75 | 8 | 8 | 9 | 33 |
| sys-casa_sub-HC011 | 2022-01-26 | 0.5 | 37 | 18 | 3 | 42 | 6 |
| sys-casa_sub-HC012 | 2022-01-26 | 0.5 | 28 | 6 | 3 | 63 | 9 |
| sys-casa_sub-HC013 | 2022-02-01 | 0.5 | 40 | 15 | 7 | 38 | 7 |
| sys-casa_sub-HC014 | 2022-02-01 | 0.5 | 45 | 16 | 9 | 30 | 24 |
| sys-casa_sub-HC015 | 2022-02-01 | 0.5 | 52 | 10 | 8 | 30 | 24 |
| sys-casa_sub-HC016 | 2022-02-25 | 0.5 | 76 | 14 | 2 | 8 | 224 |
| sys-casa_sub-HC017 | 2022-02-25 | 0.5 | 70 | 24 | 3 | 3 | 506 |
| sys-casa_sub-HC018 | 2022-02-25 | 0.5 | 62 | 18 | 4 | 16 | 770 |
| sys-casa_sub-HC019 | 2022-03-04 | 0.5 | 82 | 13 | 2 | 3 | 86 |
| sys-casa_sub-HC020 | 2022-03-04 | 0.5 | 43 | 22 | 10 | 25 | 90 |
| sys-casa_sub-HC021 | 2022-03-09 | 0.5 | 90 | 7 | 2 | 1 | 102 |
| sys-casa_sub-HC022 | 2022-03-09 | 0.5 | 91 | 4 | 1 | 4 | 226 |
| sys-casa_sub-HC023 | 2022-03-09 | 0.5 | 74 | 16 | 4 | 6 | 24 |
| sys-casa_sub-HC024 | 2022-03-11 | 0.5 | 87 | 6 | 3 | 4 | 13 |
| sys-casa_sub-HC025 | 2022-03-18 | 0.5 | 75 | 13 | 4 | 8 | 28 |
| sys-casa_sub-HC026 | 2022-03-18 | 0.5 | 82 | 8 | 2 | 8 | 16 |
| sys-casa_sub-HC027 | 2022-03-18 | 0.5 | 90 | 6 | 2 | 2 | 202 |

**Table A10:**
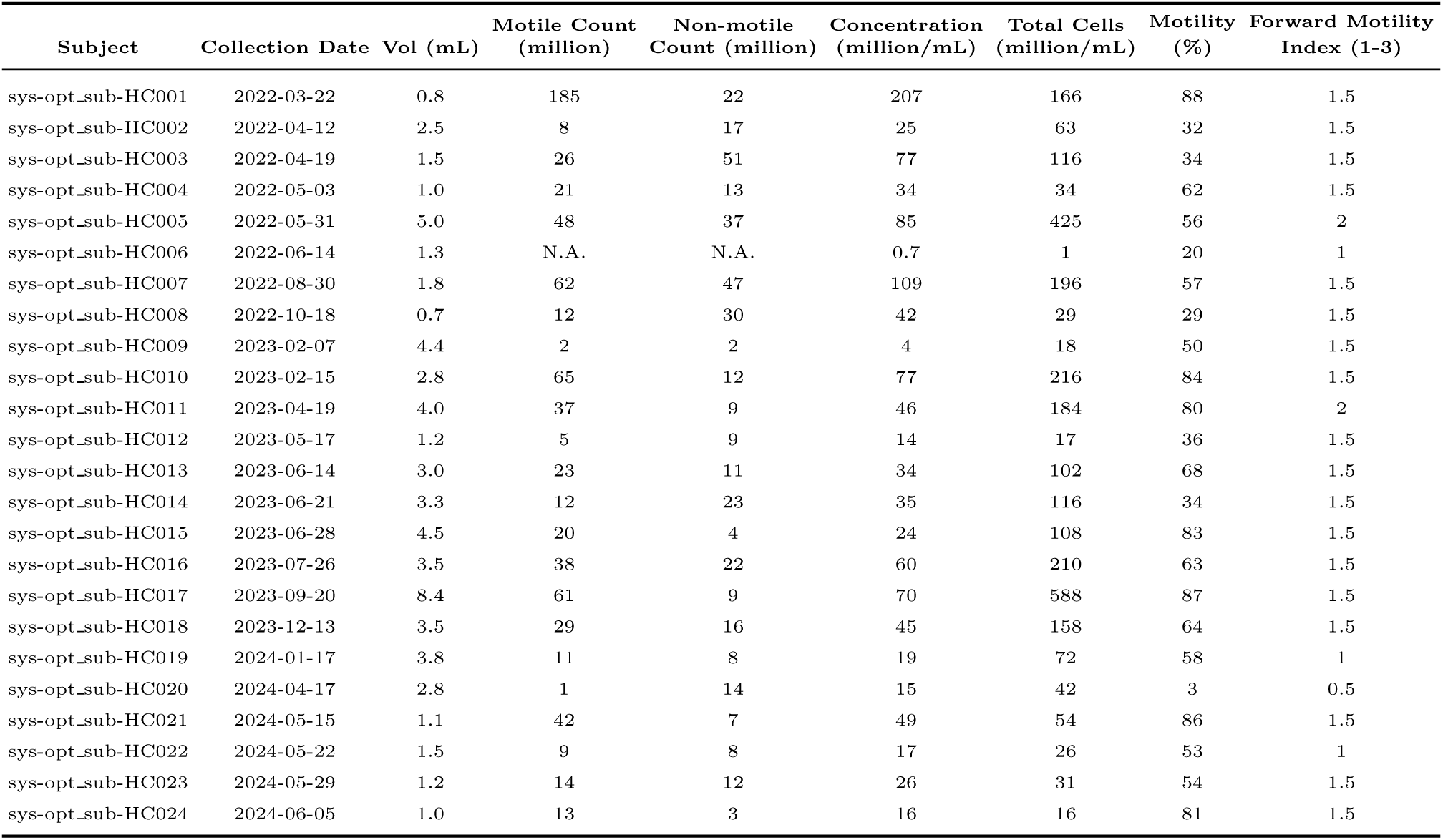
Optical Microscope Pre-Prep Motility Parameters

**Table A11:** Optical Microscope Post-Prep Motility Parameters

| Subject | Collection Date | Vol (mL) | Motile Count<br>(million) | Non-motile<br>Count (million) | Concentration<br>(million/mL) | Total Cells<br>(million/mL) | Motility<br>(%) | Forward Motility<br>Index (1-3) |
| --- | --- | --- | --- | --- | --- | --- | --- | --- |
| sys-opt_sub-HC001 | 2022-03-22 | 1.0 | 84 | 13 | 97 | 97 | 87 | 2 |
| sys-opt_sub-HC002 | 2022-04-12 | 1.0 | 7 | 12 | 19 | 19 | 36 | 1.5 |
| sys-opt_sub-HC003 | 2022-04-19 | 1.0 | 14 | 13 | 27 | 27 | 52 | 1.5 |
| sys-opt_sub-HC004 | 2022-05-03 | 1.0 | 7 | 7 | 14 | 14 | 50 | 2.5 |
| sys-opt_sub-HC005 | 2022-05-31 | 1.0 | 107 | 25 | 132 | 132 | 81 | 3 |
| sys-opt_sub-HC006 | 2022-06-14 | 0.5 | N.A. | N.A. | 2-3 | 1-1.5 | 5 | 0.5 |
| sys-opt_sub-HC007 | 2022-08-30 | 1.0 | 45 | 9 | 54 | 54 | 83 | 2.5-3.0 |
| sys-opt_sub-HC008 | 2022-10-18 | 1.0 | 4 | 6 | 10 | 10 | 40 | 1.5 |
| sys-opt_sub-HC009 | 2023-02-07 | 1.0 | 18 | 15 | 33 | 33 | 55 | 2 |
| sys-opt_sub-HC010 | 2023-02-15 | 1.0 | 108 | 15 | 123 | 123 | 88 | 2.5 |
| sys-opt_sub-HC011 | 2023-04-19 | 1.0 | 51 | 12 | 63 | 63 | 80 | 3 |
| sys-opt_sub-HC012 | 2023-05-17 | 1.0 | 2 | 2 | 4 | 4 | 50 | 1.0 |
| sys-opt_sub-HC013 | 2023-06-14 | 1.0 | 20 | 12 | 32 | 32 | 63 | 1.5 |
| sys-opt_sub-HC014 | 2023-06-21 | 1.0 | 18 | 14 | 32 | 32 | 54 | 1.5 |
| sys-opt_sub-HC015 | 2023-06-28 | 1.0 | 59 | 3 | 62 | 62 | 95 | 1.5 |
| sys-opt_sub-HC016 | 2023-07-26 | 1.0 | 60 | 42 | 102 | 102 | 58 | 1.5 |
| sys-opt_sub-HC017 | 2023-09-20 | 1.0 | 69 | 17 | 86 | 86 | 80 | 2.5 |
| sys-opt_sub-HC018 | 2023-12-13 | 1.0 | 37 | 43 | 80 | 80 | 46 | 2.0 |
| sys-opt_sub-HC019 | 2024-01-17 | 1.0 | 29 | 5 | 34 | 34 | 85 | 2 |
| sys-opt_sub-HC020 | 2024-04-17 | 0.5 | 2 | 9 | 11 | 6 | 13 | 1 |
| sys-opt_sub-HC021 | 2024-05-15 | 1.0 | 38 | 21 | 59 | 59 | 64 | 1.5 |
| sys-opt_sub-HC022 | 2024-05-22 | 1.0 | 22 | 7 | 29 | 29 | 76 | 1.5 |
| sys-opt_sub-HC023 | 2024-05-29 | 1.0 | 6 | 5 | 11 | 11 | 55 | 1.2 |
| sys-opt_sub-HC024 | 2024-06-05 | 1.0 | 13 | 3 | 16 | 16 | 81 | 1.5 |

## Notes

https://huggingface.co/datasets/DFL-KamLab/HSMV_A-Dataset-of-Human-Sperm-Microscopy-Videos

### Summary of Updates

(1) Additional sperm trajectory ground-truth information (2) Dataset name change

